# RNF213 loss of function reshapes vascular transcriptome and spliceosome leading to disrupted angiogenesis and aggravated vascular inflammatory responses

**DOI:** 10.1101/2022.01.18.476838

**Authors:** Liyin Zhang, Sherif Rashad, Yuan Zhou, Kuniyasu Niizuma, Teiji Tominaga

**Author notes:** Corresponding author details: Sherif Rashad, M.D, Ph.D., Kuniyasu Niizuma, MD, PhD.

## Abstract

**Rationale:** Moyamoya disease (MMD) is a rare cerebrovascular occlusive disease that affects Asian population more often. The pathogenesis of MMD is related to mutation in RNF213 gene. However, why, and how RNF213 mutation leads to MMD is still not fully understood.

**Objective:** Analyze the impact of RNF213 loss of function on vascular cells and the observed changes correlate with MMD.

**Methods and results:** RNF213 KD was conducted in HUVEC and vascular smooth muscle cells (vSMCs). First, HUVEC cells showed alteration of angiogenesis, migration under LPS stimulation, Leukocyte endothelial transmigration, and endothelial-to-vSMCs communication.

Transcriptome analysis revealed downregulation of genes regulating cell division and mitosis. Interestingly, Alternative splicing (AS) analysis revealed hundreds of AS events to occur after RNF213 KD in various types of AS. The alternatively spliced genes showed minimal overlap with transcriptome profiling. Pathway analysis revealed many processes and pathways to be regulated by AS events observed. Transcriptome profiling was also performed after LPS treatment and revealed the basis for increased sensitivity to LPS observed in our analysis. AS changes were also observed after LPS treatment in RNF213 KD HUVEC. Different types of AS showed different patterns of convergence or divergence in terms of the regulated pathways after LPS treatment. Finally, transcriptome and AS analysis was performed in vSMCs and showed various processes that impact vSMCs function and phenotype in RNF213 loss of function.

**Conclusion:** Our data provide a wealth of information on RNF213 gene function in vascular tissues and shed important light on different processes that contribute to MMD pathogenesis. Our results signify an immune-driven and hemodynamically linked process of MMD initiation and progression.

## Introduction

Moyamoya disease (MMD) is a rare occlusive cerebrovascular disease that has high prevalence in east Asian populations^1,2^. MMD is characterized by progressive stenosis and occlusion of the internal carotid arteries (ICAs) and the development of basal brain collaterals^1,2^. While the standard-of-care, surgical treatment, and postoperative management of surgical complications have been fairly standardized in MMD therapy^2–5^, the etiology of MMD is not well understood.

MMD is a genetic disease, and polymorphism in the gene RNF213 was identified as a susceptibility gene for its development^6^. RNF213 is an E3 ubiquitin ligase/AAA+ ATPase that belongs to RING (really interesting new gene) finger family^7^. However, the exact mechanism by which RNF213 mutations induce MMD is not known. Moreover, the rate of RNF213 mutations in healthy Japanese population ranges between 1.4∼2.7%^6,8^. This, and the association of MMD with autoimmune diseases^9^, led to the hypothesis that RNF213 mutations alone do not cause MMD, but a compendium of environmental and immune factors might be at play^7,10^. Indeed, with the advent of modern technologies that allowed the early diagnosis of MMD in its early stages, understanding the pathophysiologic mechanisms governing its development and progression would allow for potential therapies to be developed or repurposed in order to halt the disease. Previous research showed that RNF213 regulates immune function and response to bacterial infection^7,11^, involved in regulation of cytoplasmic lipid droplets^12^, regulates vascular development visa non-canonical WNT signaling pathway^13^, regulates neuromuscular development in zebrafish^14^, and regulates inflammatory responses and angiogenesis in endothelial cells^15^.

Despite recent findings regarding the function of RNF213, the impact of its mutations and their role in MMD development remain unknown. Furthermore, RNF213 is far from being understood nor it is fully characterized. MMD pathophysiology poses interesting yet challenging questions; Why does the vascular stenosis occurs only in a specific region in the terminal ICA? What environmental or immune factors might be at play in MMD? And what are the vascular functions of RNF213?

It is important to note that many of what we know about RNF213 came from research using non-vascular systems. How do previous data correlate to the role of RNF213 in vascular cells (endothelial cells (ECs), vascular smooth muscle cells (vSMCs) and others) is yet to be evaluated.

In this work, we evaluated the impact of RNF213 knockdown (KD) on ECs function and behavior as well as on vSMCs. Using RNA-sequencing we performed in-depth transcriptional and epigenetic profiling of the impact of RNF213 KD (simulating loss of function of RNF213 as reported in its mutation) on ECs and vSMCs biology as well as EC-to-vSMCs and EC-to-immune cells interactions. Our results support an immune exacerbated vascular dysfunction in RNF213 KD that can explain certain aspects of MMD pathophysiology.

## Methods

### Cell culture and reagents

Human umbilical vein endothelial cells (HUVEC) were purchased from ATCC (Cat# CRL-1730) and Primary aortic smooth muscle cells (vSMCs) were purchased from ATCC (Cat# PCS-100-012) and cultured in Medium 199 (Cat# 31100-035; Invitrogen Corporation, Carlsbad, CA), 10μg Fibroblast Growth Factor (FGF) (Wako Cat# 064-04541), 20% fetal bovine serum (Cat# FB- 1360/500; JRH Biosciences, Lenexa, KS), and 1% penicillin/streptomycin (Cat# 15140122; Invitrogen) on 1% gelatin coated dishes and flasks. For quiescent medium of SMCs, D-MEM/F-12 (Thermo Fisher 12500062), 10ml/1L of Insulin-Transferrin-Selenium-Ethanolamine (ITS -X; Gibco Cat# 51500056) and 1% penicillin/streptomycin were used.

Human macrophage cell line (MV-4-11) was purchased from ATCC (Cat# CRL-9591) and cultured in IMDM (ATCC Cat# 30-2005), 10%FBS, and 1% penicillin/streptomycin.

### siRNA treatment

RNF213 siRNA was purchased from Thermo Fischer (Silencer select siRNA, Cat# s33658) Control (Mock) siRNA was purchased from Thermo Fischer (Silencer select siRNA, Cat# 4390843)

siRNA knockdown (KD) of RNF213 was achieved by platting HUVEC or vSMCs in serum free M199 medium and transfecting the cells using Lipofectamine RNAiMAX (Thermo Fischer Cat# 13778150). Cells were incubated for 48 hours before changing the medium for full growth medium (GM). Cells were allowed to rest for 24 hours before any experiments or analysis. KD efficiency was confirmed using qPCR as good antibodies against RNF213 for western blotting are not readily available.

### Wall shear stress (WSS) analysis

Endothelial cell response to WSS changes was performed as published previously^16^. In-short, HUVEC cells were cultured until confluent on the bottom surface of a ϕ35-mm plastic dish, with a proliferation medium (PM) containing Medium 199 (Invitrogen, Cat#31100-035), 20% fetal bovine serum (JRH Biosciences, Cat#FB-1360/500), and 1% penicillin/streptomycin (Invitrogen, Cat #15140122). A flow loop was constructed by connecting a pulse damper, flow chambers, a reservoir, and a roller pump with silicone tubes. Cells were exposed to different magnitudes of WSS (0.4, 2, and 10 Pa) in parallel plate flow chambers (PPFCs) for 24 hours using PM at 37°C in an atmosphere of 95% air/5% CO2. The selection of these values was based on previous *in vivo* doppler ultrasound measurements^17^. Cells were collected by scrapping and lysed in TRIzol reagent. More details on the flow chamber design, characterization, and parameters can be found in our previous report^16^.

### RNA extraction

Cells were lysed in TRIzol reagent (Qiazol, Qiagen, Cat#79306). RNA was extracted using the miRNeasy mini kit (Qiagen, Cat# 217004) with DNase digestion step. RNA concentration and purity was analyzed using nanodrop One. RNA integrity number (RIN) was analyzed using Agilent Bioanalyzed RNA 6000 nano (Cat# 5067-1511)

### Real-time PCR (qPCR) analysis

cDNA was synthesized from RNA using Superscript III first strand synthesis system (Invitrogen, Cat# 18080051). qPCR was performed using GoTaq qPCR Master Mix (Promega, Cat# A6102) on CFX96 thermal cycler (Bio-Rad). qPCR was performed using GoTaq qPCR Master Mix (Promega, Cat# A6102) on CFX96 thermal cycler (Bio-Rad). qPCR was conducted with 4 replicates per group. qPCR was analyzed using the ΔΔCT method with Actin-β as an internal control.

Primers used in this study:

RNF213: F: 5’TCGTAACAACACGGAAGTGGAGA, R: 5’AGCAGAGTGCAGGTCGATGAA
Actin-β: F: 5’CATGTACGTTGCTATCCAGGC, R: 5’CTCCTTAATGTCACGCACGAT
ICAM1: F: 5’TGTATGAACTGAGCAATGTGCAAGA, R: 5’CACCTGGCAGCGTAGGGTAA
SELE: F: 5’CACTCAAGGGCAGTGGACACA, R: 5’CAGCTGGACCCATAACGGAAAC
PECAM1: F: 5’CAGAGTACCAGGTGTTGGTGGAA, R: 5’GACAGAACAGTTGACCCTCACGA
SELP: F: 5’GAAGATGGTCAGCTACTCCACCA, R: 5’GTTCCCGGATGATGCCTACAG
VCAM1: F: 5’CGAAAGGCCCGTTGAAGGA, R: 5’GAGCACGAGAAGCTCAGGAGAAA

### RNA sequencing

mRNA sequencing was conducted using NEBNext Ultra II Directional RNA Library Prep kit (Cat# E7765S) as per manufacturer’s instructions. All samples using had RIN > 9. Quality control of libraries was performed using Agilent DNA 1000 kit (Cat# 5067-1504). NEBNext Library Quant Kit (Cat# E7630L) was used for quantifying libraries. Libraries were pooled and sequenced on Hiseq-X ten (150bp, paired end) by Macrogen Japan. All sequencing experiments were performed with 3 replicates per group.

### RNA sequencing data analysis

Quality control for Raw fastq files was performed using FastQC. Reads were trimmed using Trimmomatic^18^ and adaptor sequences and low-quality reads removed. Reads were aligned to human reference genome (Hg38 assembly from UCSC) using Hisat2^19^ with splice aware options using GTF file for reporting. Read counting and transcript assembly was performed using Stringtie^20^, and differential gene expression was performed using DESeq2^21^. Fold change > 1.5 (Fold change > 2 was set for the LPS experiment) plus FDR < 0.05 were criteria for statistical significance of differentially expression genes (DEGs).

Alternative splice (AS) analysis was performed using the rMATs-turbo suite^22^. Statistical significance of alternative splicing events was considered when FDR < 0.05 and delta-Psi (ΔΨ) > 0.2. Sashimi plots were created using rmats2sashimiplot (https://github.com/Xinglab/rmats2sashimiplot/issues/72)

Gene ontology and pathway analysis was performed using Metascape^23^. Only genes satisfying the selection criteria in DEGs analysis or AS analysis were selected for pathway analysis.

### Tube-formation assay

24- well plate was coated with Geltrex (Gibco Cat# A1413202) for tube formation assay of HUVEC. 35000 cells were seed after siRNA knockdown, and cells observed for 2∼ 24 hours with phase contrast microscope

### Migration (scratch) assay

After siRNA to knockdown RNF-213, the HUVEC cells was reseed into 6-well plate with 130000 cells with or without 1μg/ml LPS overnight to reach to confluency. A sterile 1000μl pipette tip was used to make a scratch in the midline of each well and observation was done at 0h, 4h, 6h and 24h under electron microscope. A small cross scratch was also performed to mark the location of photography for consistency in the measurements.

### Endothelial-vascular smooth muscle cells co-culture system

ϕ35mm silicon ring were put into the 35mm dish, SMCs imbedded in Atelocollagen acidic solution (IPC-50) were seeded in the middle part of the ring and cultured in proliferation medium: M199, 10μg FGF, 20% fetal bovine serum, and 1% penicillin/streptomycin until it reached confluency. Medium was then changed into D-MEM/F-12, Insulin-Transferrin-Selenium-X, and 1% penicillin/streptomycin for 7 days. siRNA transfected HUVECs were seeded on fibronectin coated 5μm SMWP membrane filters (Merck Millipore Cat# SMWP02500) and inserted atop the silicon ring [see figure 7].

### Leukocyte transmigration assay

For transmigration assay, CytoSelect Leukocyte Transmigration Assay (CBO Cat# CBA-212) was used. After siRNA transfection, 10^5^ HUVEC cells were seeded on the upper chamber of the transmigration plate for 24h with or without LPS treatment (100ng/ml). MV-4-11 macrophages were added after 24h, in the upper chamber for 4h. The transmigrated macrophages in the lower chamber were then observed using fluorescence plate reader as per manufacturer’s instruction.

## Results

➢ **RNF213 is a mechano-responsive gene that regulates endothelial angiogenesis** A hallmark of MMD is the terminal progressive ICA stenosis followed by the formation of inadequate collaterals at the skull base^2^. Nonetheless, the factors initiating this stenosis are not known. Vascular pathologies, in general, are caused by a wide array of factors interacting with the vascular wall leading to the initiation or progression of said pathologies. One of the most important factors is the interaction between blood hemodynamics and endothelial cells (ECs)^24^. This interaction causes and modulates many vascular pathologies such as stenosis, aneurysms, and atherosclerosis^16,24,25^. We have previously shown that MMD symptomatology is impacted by complex hemodynamics arising from the complex morphology of the MMD associated ICA stenosis in early stage MMD^26^. However, our previous work does not provide a causal link between vascular hemodynamics and MMD initiation, but it was rather focused on the impact of MMD induced ICA stenosis on the hemodynamics leading to transient ischemic attacks (TIA). To that end, we used our previously published system for interrogating flow stress impact on ECs to study the effect of wall shear stress (WSS) on RNF213 gene expression^16^. Our analysis revealed that RNF213 is a mechanoresponsive gene, as it was significantly upregulated in response to high WSS (10 pascals) [Figure 1A]. Importantly, the same WSS value was associated with a pro-angiogenic response in ECs in our previous work^16^, indicating that RNF213 might play a role in the blood flow shear stress induced angiogenesis. This is in line with the previously reported role of RNF213 on regulating angiogenesis^6^. **Figure 1:**
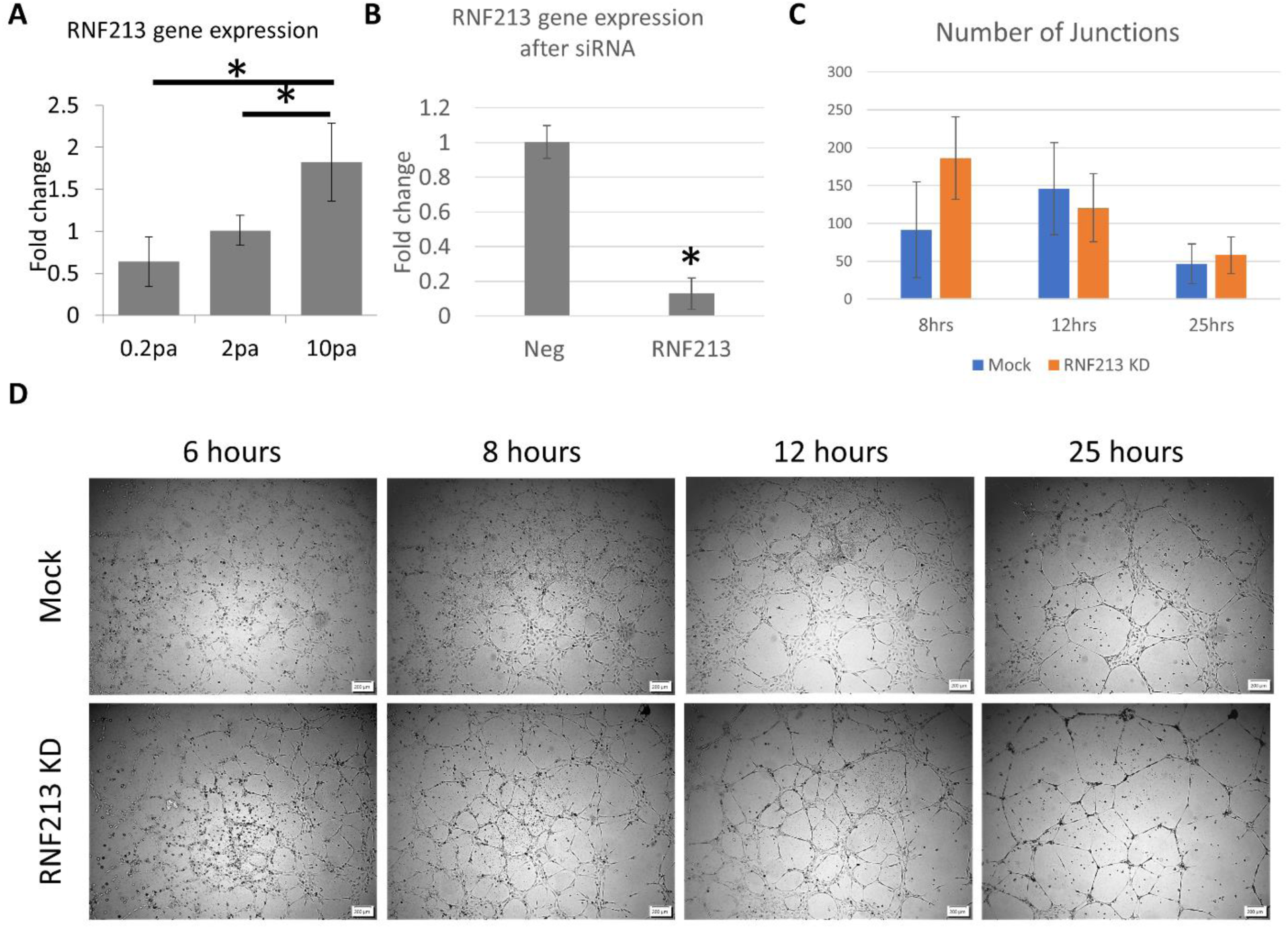
RNF213 is a mechano-sensitive gene that alters endothelial angiogenesis. A: RNF213 gene expression analysis using qPCR after exposure to different levels of WSS. B: siRNA KD of RNF213 in HUVEC cells yielded > 80% reduction of RNF213 gene expression. C: Analysis of the number of junctions in the tubes formed by HUVEC cells transfected by RNF213 siRNA or Mock siRNA. D: Phase contrast images from tube formation assay in RNF213 of Mock transfected HUVEC cells. Note the thicker tube walls in Mock cells compared to RNF213 cells. Also, at 6 hours, RNF213 transfected cells formed robust tubes compared to the immature tubes formed in the Mock cells. Asterisk: Statistically significant (*p* < 0.05, Fold change > 1.5). To test the impact of RNF213 on vascular angiogenesis, we used siRNA-mediated knockdown (KD) and tube formation assay. Our protocol led to >80% KD in RNF213 gene expression [Figure 1B]. Tube formation assay revealed interesting pattern [Figure 1C and D]. First, RNF213 KD cells formed mature tubes earlier than Mock cells [6 hours in Figure 1D]. Moreover, the walls of the tubes in Mock cells were evidently thicker than RNF213 KD cells. This can be reflective of the weak collaterals formed in MMD patients (i.e., the Moyamoya vessels at the skull base), which are characterized by being weak and inefficient and prone to rupture^4^.
➢ **RNF213 KD alters endothelial transcriptome and epigenome** To further understand the influence of RNF213 loss of function on vascular biology, we conducted RNA sequencing (RNA-seq) analysis on ECs after RNF213 siRNA. Transcriptome profiling of RNF213 function using *in vitro* systems was surprisingly lacking in the literature, thus, a holistic understanding of RNF213 role in ECs cannot be achieved. Our RNFA-seq analysis revealed hundreds of genes to be up and down regulated after RNF213 KD [Fold change > 1.5, FDR < 0.05. Figure 2A]. We analyzed the gene ontology (GO) enrichment and gene set enrichment (GSE) for the differentially expressed genes (DEGs) using Metascape^23^. GO Biological processes term enrichment revealed an interesting pattern of clustering, where many processes related to cell division and proliferation were downregulated in RNF213 KD [Figure 2B]. Cluster analysis revealed the existence of 2 major clusters, one related to cell division/proliferation, which was exclusively downregulated in RNF213 KD cells, and another cluster related to organellar organization and cell motility, which was shared enriched in up and downregulated genes [Figure 2C, supplementary figure 1A]. GO molecular functions analysis revealed the upregulation of Integrin binding after RNF213 KD, which might regulate cellular responses to flow stress^27^, as well as other terms that reflect changes in cellular motility, cell surface receptors, or interaction with other cells [Supplementary figure 1B]. GO cellular components analysis revealed the top 2 enriched terms to be related to DNA and cell division, and both to be downregulated in RNF213 KD cells [Figure 2D]. Cluster analysis revealed 2 main clusters; one related to cell division/proliferation and was downregulated in RNF213 KD, and another related to cytoplasmic and exocytic vesicles and was upregulated in RNF213 KD cells [Supplementary figure 2]. **Figure 2:**
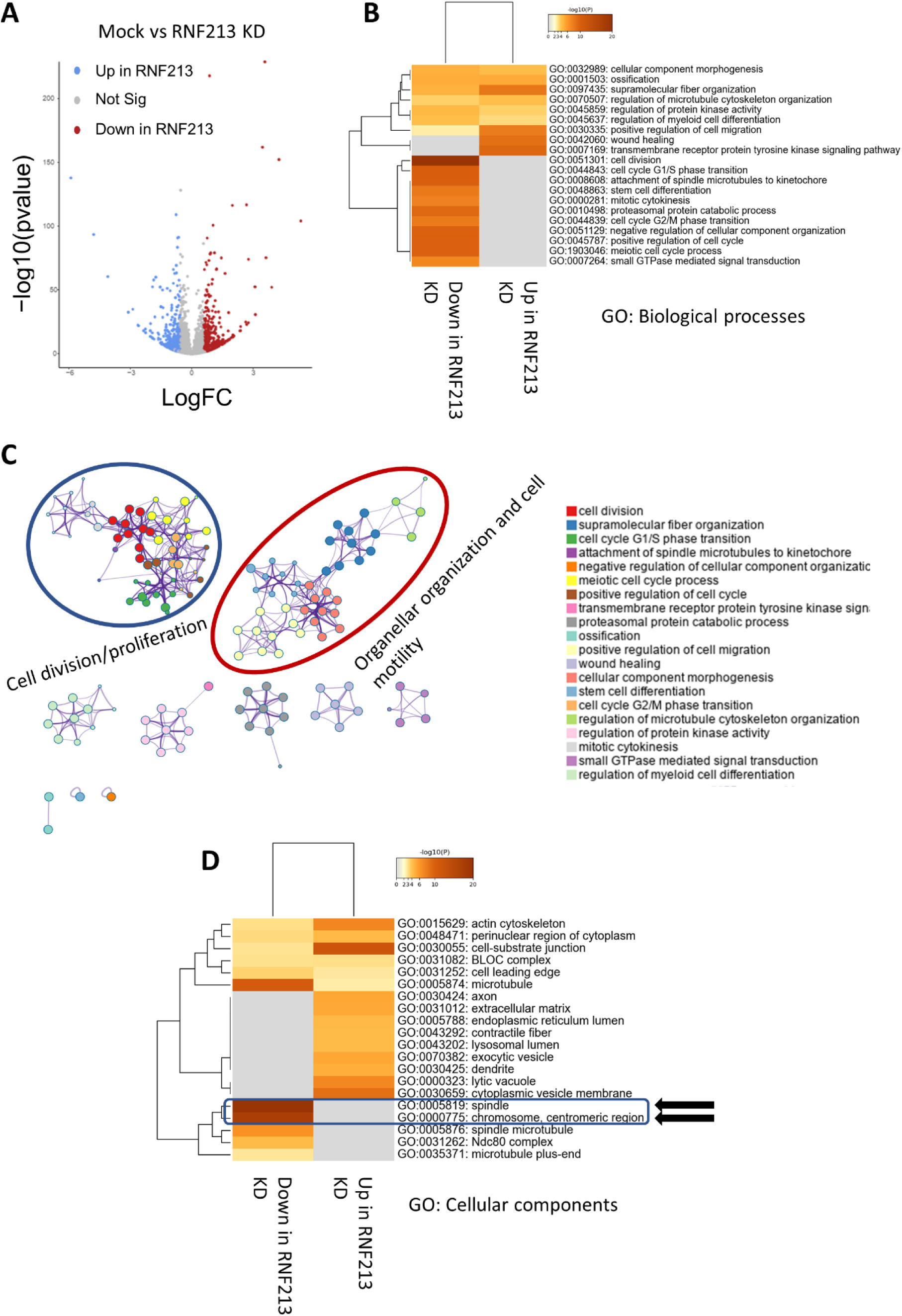
Transcriptome profiling after RNF213 KD in HUVEC. A: Volcano plot of differentially expressed genes after RNF213 KD. B: GO biological processes (BP) analysis revealing downregulation of genes related to cell division and proliferation after RNF213 KD. C: Cluster analysis of GO BP enriched terms showing two main clusters related to Cell division/proliferation and organellar organization and cell motility. D: GO cellular components (CC) enriched terms showing significant enrichment of terms linked to cell division and chromosome replication in the downregulated genes after RNF213 KD. Protein-Protein interaction (PPI) analysis further supported the observed impact of RNF213 on cellular transcriptome. Clusters related to cell division, ubiquitin activity, and mitochondria were downregulated in RNF213 [Supplementary figure 3A]. While clusters related to vesicle transport and cell adhesion being upregulated after RNF213 KD [Supplementary figure 3B]. PPI analysis using both gene sets (up and downregulated) revealed clusters related to angiogenesis as well as other processes related to vascular endothelial cell division and organellar organization [Supplementary figure 4, supplementary table 1]. Alternative splicing (AS) is well known mechanism that regulates cellular proteome diversity, protein translation, mRNA stability and mRNA expression as well as being a part of the oxidative stress response^28–30^. We interrogated our RNA-seq data to evaluate whether RNF213 KD could impact AS machinery or not. Several methods exist to analyze AS, we utilized local splice variant (LSV) analysis, defined as a split in a splice graph into or from a single exon, termed the reference exon, which can reveal layers of complexity in the AS process^22,31^. LSV analysis revealed hundreds of AS events in 5 AS types: Skipped exon (SE), retained intron (RI), mutually exclusive exon (MXE), alternative 5’ splice site (A5SS), and alternative 3’ splice site (A3SS) [Figure 3, supplementary figure 5]. Interestingly, there was nearly no overlap between genes that showed LSV events enrichment and DEGs [Supplementary figure 6], indicating that LSV changes induced by RNF213 loss of function are another layer of gene regulation separate from transcriptome changes. SE events were the highest in terms of number of significant events, followed by A3SS, A5ss, and MXE. RI events were the lowest in terms of numbers [Figure 3B]. GO analysis revealed some overlap between various AS types, however, many unique pathways were also regulated by each AS type [Figure 3C]. However, in-depth GO analysis revealed that each LSV event type regulated different processes or pathways [Supplementary figure 7]. For example, top enriched pathways in SE events were linked to DNA damage pathways and protein quality control processes, while in A5SS the top term was cytoplasmic translation [Supplementary figure 7]. Indeed, each AS type could impact mRNA translation or stability differentially leading to the generation of different protein isoforms [in the case of SE or MXE] or introduction of early stop codons, introns, and non-sense mediated decay of mRNAs [A5SS, A3SS, and RI]^32^. The specificity of each AS type to different processes indicate complex regulation and multilayered impact of RNF213 loss of function on endothelial homeostasis. Future work should aim to identify and interrogate these mechanisms in the context of RNF213 biology and MMD pathophysiology. **Figure 3:**
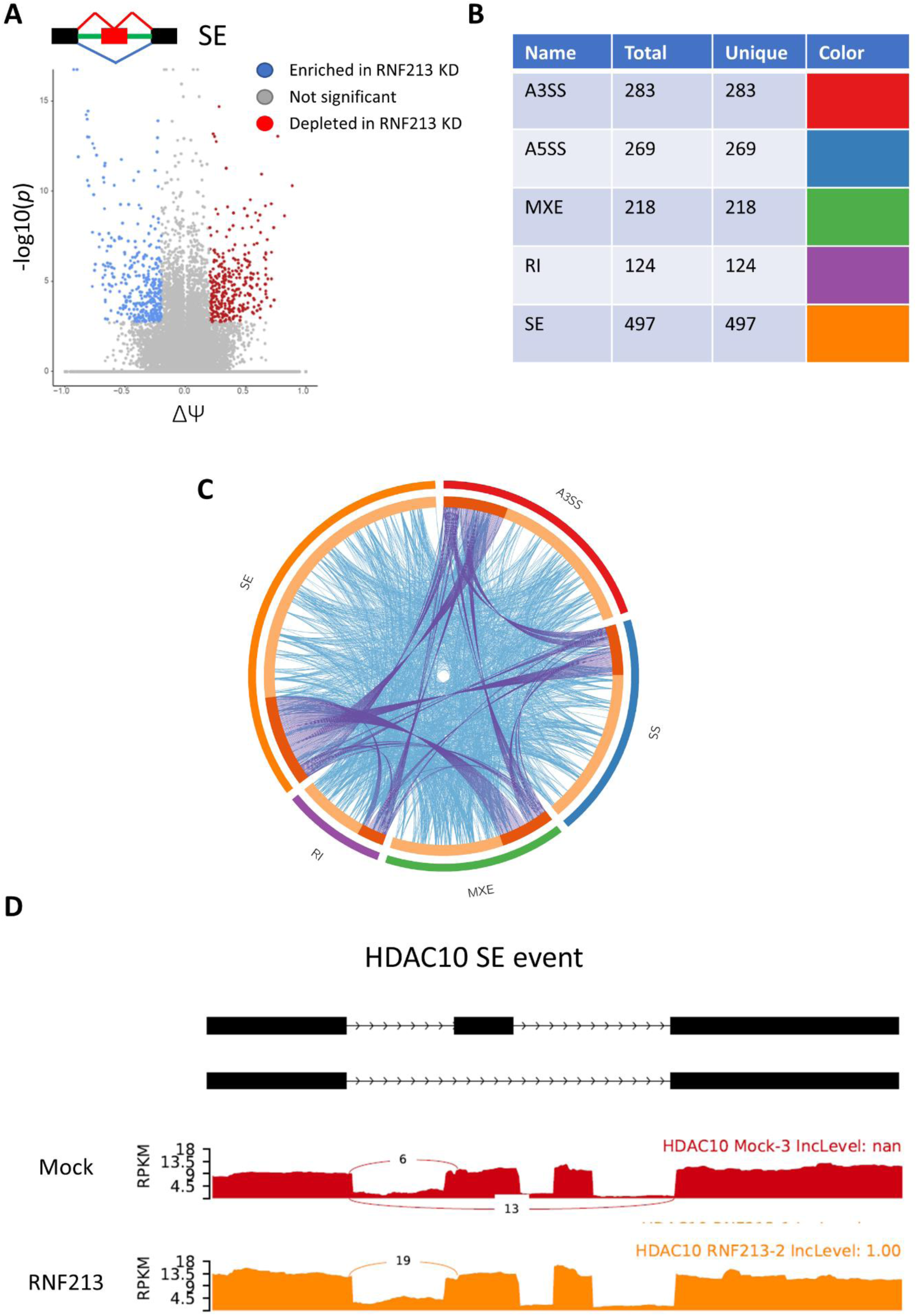
RNF213 KD in HUVEC leads to significant changes at the level of mRNA splicing. A: Volcano plot of skipped exon (SE) events after RNF213 KD. (ΔΨ = Delta psi). B: Table showing the number of events pertaining to each LSV subtype. Total genes = genes in input for each AS subtype. Unique = genes unique to each AS subtype. C: Circos plot showing overlap of alternatively spliced genes (purple curved lines) and enriched GO terms (light blue lines) between different LSV subtypes. D: Example of AS after RNF213 KD: Sashimi plot of skipped exon event in HDAC10.
➢ **RNF123 regulates endothelial response to LPS and leukocyte transmigration** RNF213 is reported to regulated ECs response to inflammatory stimuli^15^. We performed qPCR analysis after exposing wild-type ECs to lipopolysaccharide (LPS, 100ng/ml) for 24 hours and observed strong upregulation of RNF213 gene expression [Figure 4A]. To examine potential impact of RNF213 loss of function on ECs behavior during inflammation, we performed migration assay (wound scratch test) in the presence or absence of LPS (1μg/ml). RNF213 KD strongly aggravated the LPS induced retardation of endothelial migration [Figure 4B, supplementary figure 8]. **Figure 4:**
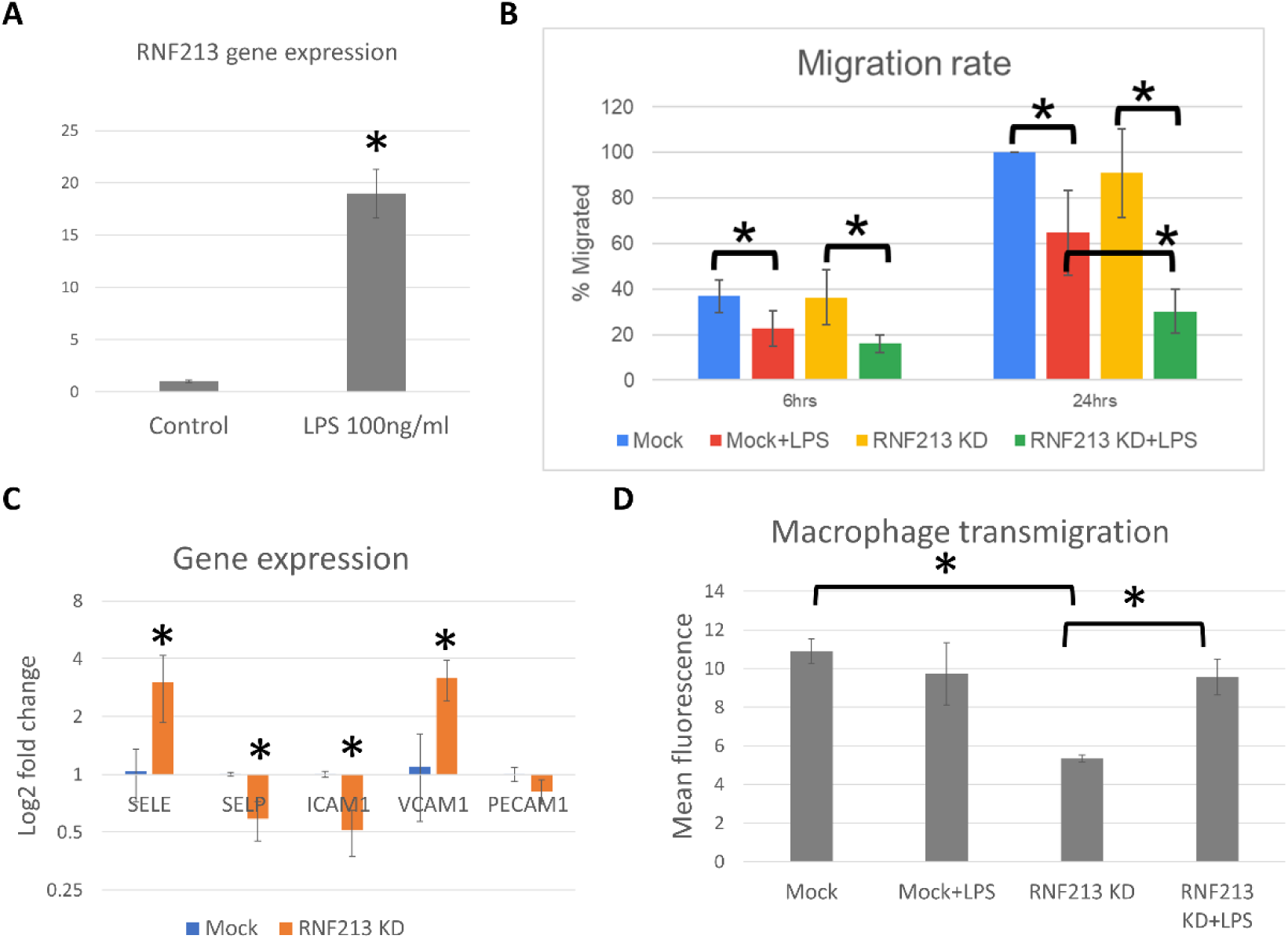
RNF213 regulates endothelial responses to LPS. A: qPCR analysis of RNF213 gene expression levels after low dose (100ng/ml) LPS stimulation for 24 hours. B: Endothelial migration assay (scratch/wound assay) showing significant reduction in HUVEC migration rates after LPS treatment (1ug/ml) and RNF213 siRNA. C: qPCR analysis of gene expression levels of cell surface receptors that regulate leukocyte adhesion and transmigration after RNF213 KD. D: Leukocyte transmigration assay showing reduction of Leukocyte transmigration after RNF213 KD, with increased sensitivity after low dose LPS (100ng/ml) evident by increased transmigration. These results confirm the role of RNF213 in regulating endothelial inflammatory responses. Given this notion, we interrogated another aspect of endothelial function under inflammation; endothelial-leukocyte interaction and leukocyte transmigration. During inflammation, leukocytes interact with endothelial cells’ surface receptors to initiate the process of transmigration towards the site of inflammation^33–35^. We began by interrogating varies EC surface receptors involved in the process of leukocyte adhesion, rolling, and eventually transmigration. qPCR analysis after RNF213 KD in ECs revealed upregulation of Selectin-E (SELE) and VCAM1 [Figure 4C], while Selectin-P (SELP) and ICAM-1 were downregulated [Figure 4C]. VCAM-1 is associated with leukocyte capture and tethering as well as with rolling^34^. SELP and SELE are both associated with leukocyte rolling^34^. ICAM-1 is integral to the process of tight adhesion between Leukocytes and ECs, and the eventual transmigration^33,34^. To further evaluate the actual impact of the changes in EC surface receptors gene expression levels, we conducted a leukocyte transmigration assay. Macrophages were incubated in a trans-well system with Mock transfected or RNF213 KD HUVEC cells with or without low dose (100ng/ml) LPS stimulation. RNF213 knockdown was associated with lower transmigration as compared to Mock transfected cells, in agreement with the observed downregulation of ICAM-1 [Figure 4D]. What stood out is the heightened sensitivity of RNF213 KD cells to low dose LPS, evident from the increased rate of Leukocyte transmigration observed, contrary to the Mock transfected cells that did not show any change in response to the used LPS dose [Figure 4D].
➢ **Transcriptional and epigenetic impact of RNF213 KD on Endothelial response to LPS** The abovementioned data indicates that RNF213 KD exacerbates the endothelial response to LPS. To understand how this sensitivity is regulated, we performed RNA-seq after Mock or RNF213 KD cells were exposed to 1μg/ml LPS for 24 hours. First, we wanted to evaluate whether the cellular response to LPS is completely divergent in terms of how cells are responding or is it a matter of sensitivity to LPS as we hypothesized. To do so, we started by comparing the LPS treated groups with their untreated counterparts (i.e., Mock (untreated) vs Mock-LPS and RNF213 (untreated) vs RNF213 LPS). Indeed, in each comparison there were thousands of DEGs [Figure 5A & B]. Gene ontology analysis revealed strong overlap between enriched pathways and terms when we used the same comparison parameters (Mock LPS vs Mock and RNF213 LPS vs RNF213), and in both comparisons, there was an upregulation of pathways linked to cytokine production and response to infection and inflammation and downregulation of pathways related to cell division [Figure 5C]. This confirmed that the response to LPS is qualitatively similar after RNF213 KD to Mock transfected cells. The difference thus lies in the enhanced sensitivity of the cells and the exaggerated response. To confirm this hypothesis, we performed to comparisons. First, we analyzed the DEGs in the above comparisons (Mock LPS vs Mock and RNF213 LPS vs RNF213) to identify uniquely up and downregulated genes after LPS treatment in RNF213 KD and Mock transfected cells. Our analysis yielded 794 and 1037 genes uniquely up and downregulated in the Mock-LPS vs Mock comparison and 605 and 731 uniquely up and downregulated genes in the RNF213-LPS vs RNF213 comparison [Supplementary figure 9A and B]. Next, we analyzed GO enrichment in each of these unique gene datasets. The analysis revealed that while the unique genes upregulated in the RNF213 + LPS group showed strong enrichment for terms related to viral and inflammatory response [Supplementary figure 9C], the Mock + LPS group did not show such specific pattern and generally had weaker enrichment for inflammation-related terms [Supplementary figure 9D]. In the downregulated gene datasets, both RNF + LPS [Supplementary figure 10A] and Mock + LPS [Supplementary figure 10B] showed similar patterns of downregulation of cell proliferation related terms, albeit the scores were higher in the Mock + LPS group. **Figure 5:**
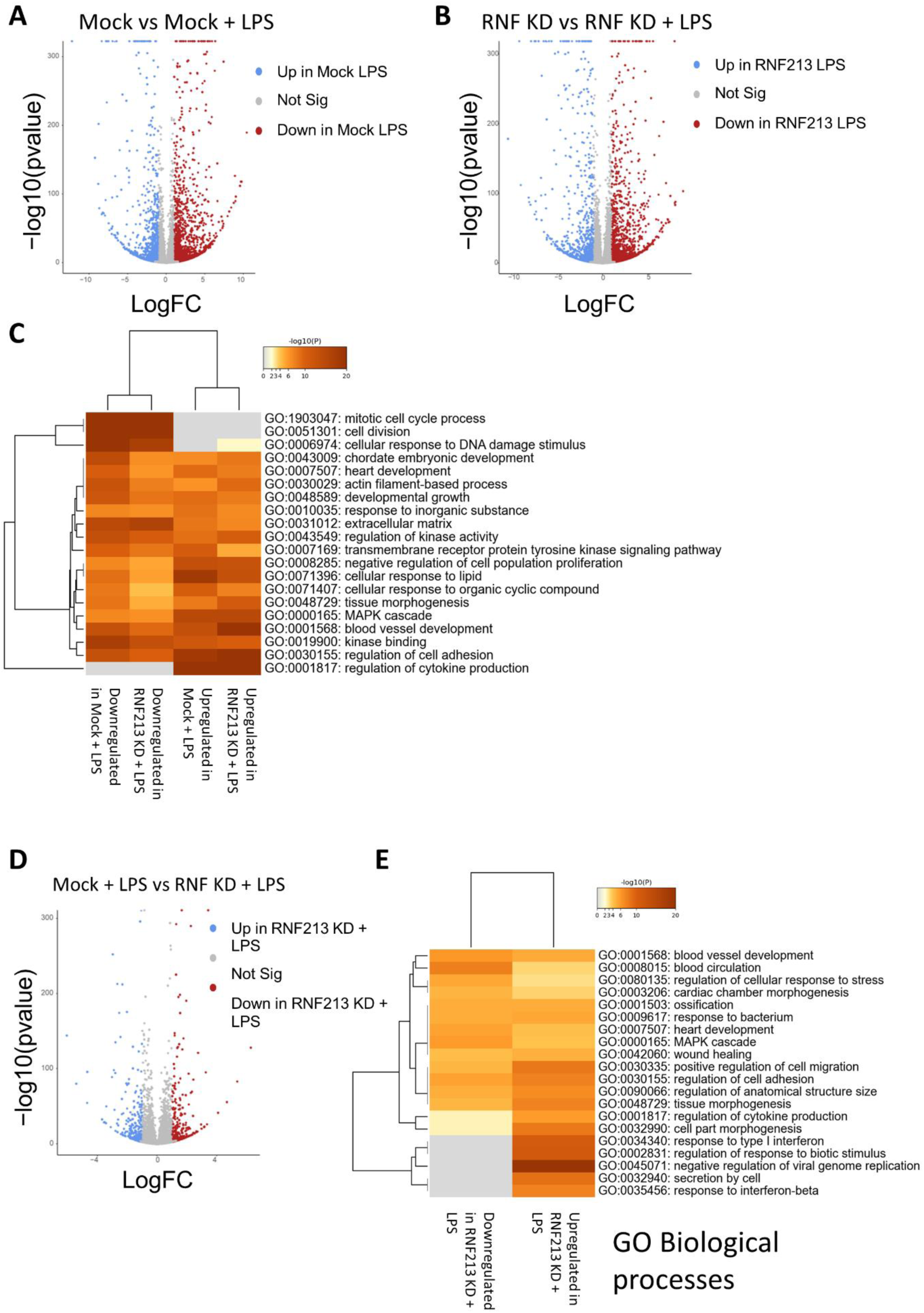
RNF213 KD alters transcriptional response to LPS. A: Volcano plot of DEGs after LPS treatment (1ug/ml) in Mock transfected cells. B: Volcano plot of DEGs after LPS treatment in RNF213 KD cells. C: GO analysis (3 terms: Biological processes, Cellular components, and Molecular functions) of enriched pathways after LPS in the comparisons shown in A and B. D: Volcano plot of DEGs after comparing LPS treated Mock and RNF213 KD cells. E: GO BP of enriched pathways in the Mock-LPS vs RNF-LPS dataset comparison. Next, we compared RNF213 KD + LPS versus Mock + LPS directly. This comparison yielded hundreds of DEGs [Figure 5D]. Looking into GO enriched pathways, it became apparent that the differentially enriched terms can faithfully explain the observed phenotypes [Figure 5E and supplementary figure 11 and 12]. RNF213 KD led to upregulation of pathways linked to inflammatory response such as pathways linked to viral infection and interferon response [Figure 5D, supplementary figure 12]. This can explain the slowed proliferation of HUVEC under LPS stimulation after RNF213 KD. It also explains the observed leukocyte transmigration enhancement at doses that did not lead to such response in Mock transfected cells. LPS treatment was previously shown to alter mRNA alternative splicing in endothelial cells^36^. Given the strong link observed between RNF213 KD and AS, we set to examine whether RNF213 KD can alter the AS landscape following LPS treatment. LPS treatment led to robust transcriptome wide AS changes after LPS treatment in HUVEC in both Mock transfected cells and with RNF213 KD, when compared with their untreated counterparts, with hundreds of events in hundreds of mRNAs observed under different AS subtypes in each group [Figure 6A and B]. Next, we examined the overlap in alternatively spliced genes/mRNA between these two datasets. A striking minimal overlap was observed [Supplementary figure 13], indicating that while LPS induced AS changes in both groups, these changes were vastly different between RNF213 KD and Mock cells. GO and pathway analysis revealed many pathways to be differentially regulated in each dataset. Looking at the top enriched pathways, it was apparent that there is a strong overlap between both groups in the enriched GO terms for the SE and MXE events [Figure 6C, Supplementary figure 14A]. RI, A5SS, and A3SS events showed more divergence in the top enriched terms [Figure 6D, Supplementary figure 14B and C]. Given that each AS type will impact the mRNAs, and in turn the regulated pathways, differently, it’s imperative, as well as interesting, to systematically analyze the outcomes of these phenomena in future work. Overall, our data indicate that RNF213 regulate endothelial response to LPS-mediated inflammation via various transcriptional as well as epigenetic mechanisms. In-depth understanding of these processes will be central to our understanding of MMD initiation and progression. **Figure 6:**
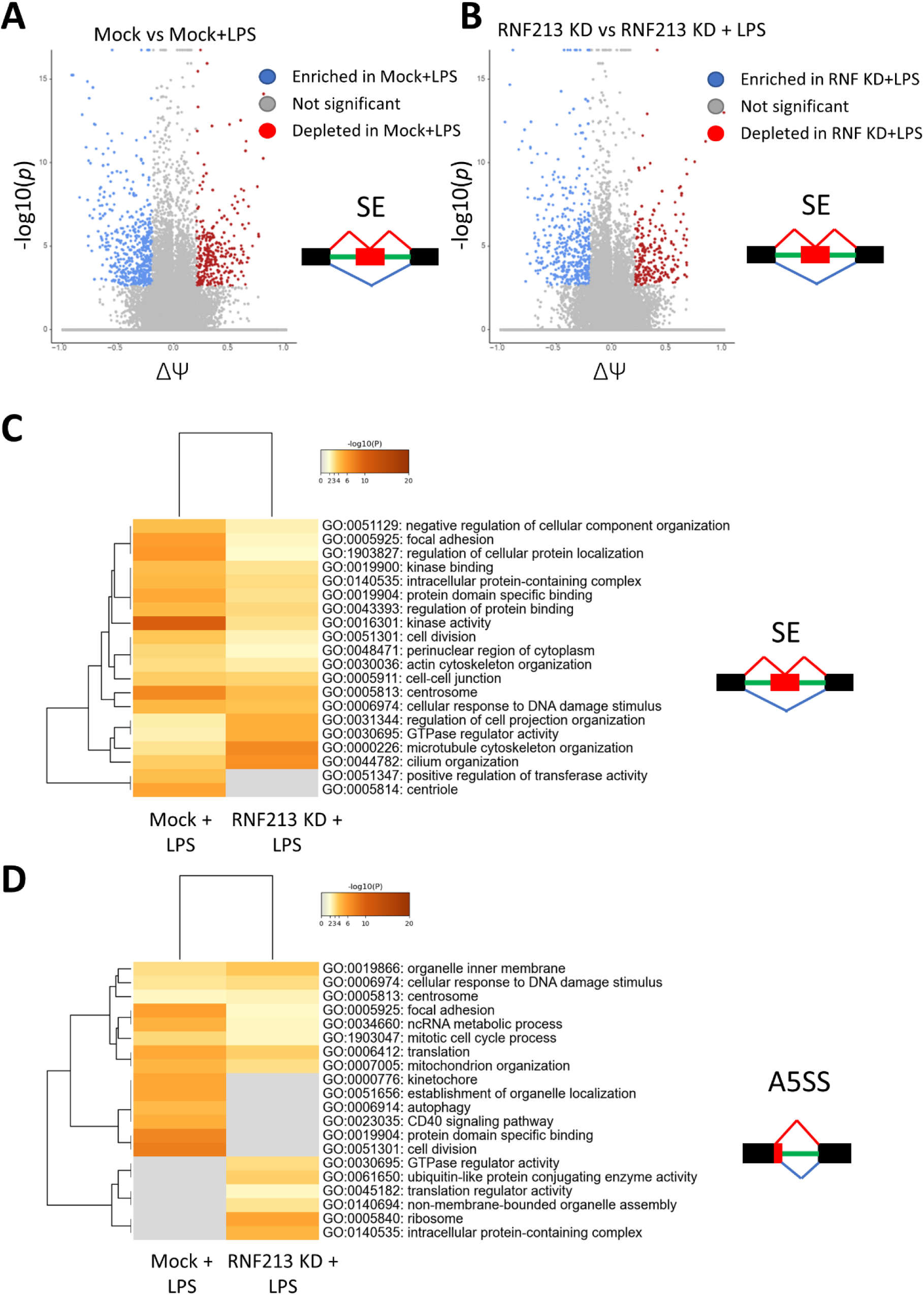
RNF213 KD alters mRNA splicing changes following LPS treatment in endothelial cells. A: Volcano plot of SE events in Mock cells before and after LPS treatment. B: Volcano plot of SE events in RNF213 KD cells before and after LPS treatment. C: GO analysis (3 terms) of SE events in the Mock or RNF213 KD groups after LPS treatment (same comparisons as in A and B). D: GO analysis (3 terms) of A5SS events in Mock or RNF213 KD groups after LPS treatment.
➢ **RNF123 regulates endothelial-to-vascular smooth muscle cells communication** ECs are known to impact vascular smooth muscle cells (vSMCs) biology through a variety of mechanisms^37,38^. Indeed, the vascular pathology of MMD extends beyond ECs, and thus we sought to identify whether RNF213 could disrupt ECs-to-vSMCs communication. We induced RNF213 KD in ECs and the co-cultured these ECs with vSMCs, followed by RNA-seq analysis of the vSMCs [Figure 7A]. Few genes were differentially expressed in vSMCs following incubation with RNF213 KD or Mock ECs [Figure 7B], however, GO analysis revealed several important pathways and processes that were impacted in vSMCs which can affect vSMCs proliferation or molecular landscape [Figure 7C]. This clearly indicates that RNF213 impacts ECs-to-vSMCs communication and would have an effect on the vascular wall and its remodulation that can explain the initiation of ICA stenosis in MMD. Nonetheless, more work is needed to validate this observation and to provide mechanistic insight to its causal factors and downstream effectors. **Figure 7:**
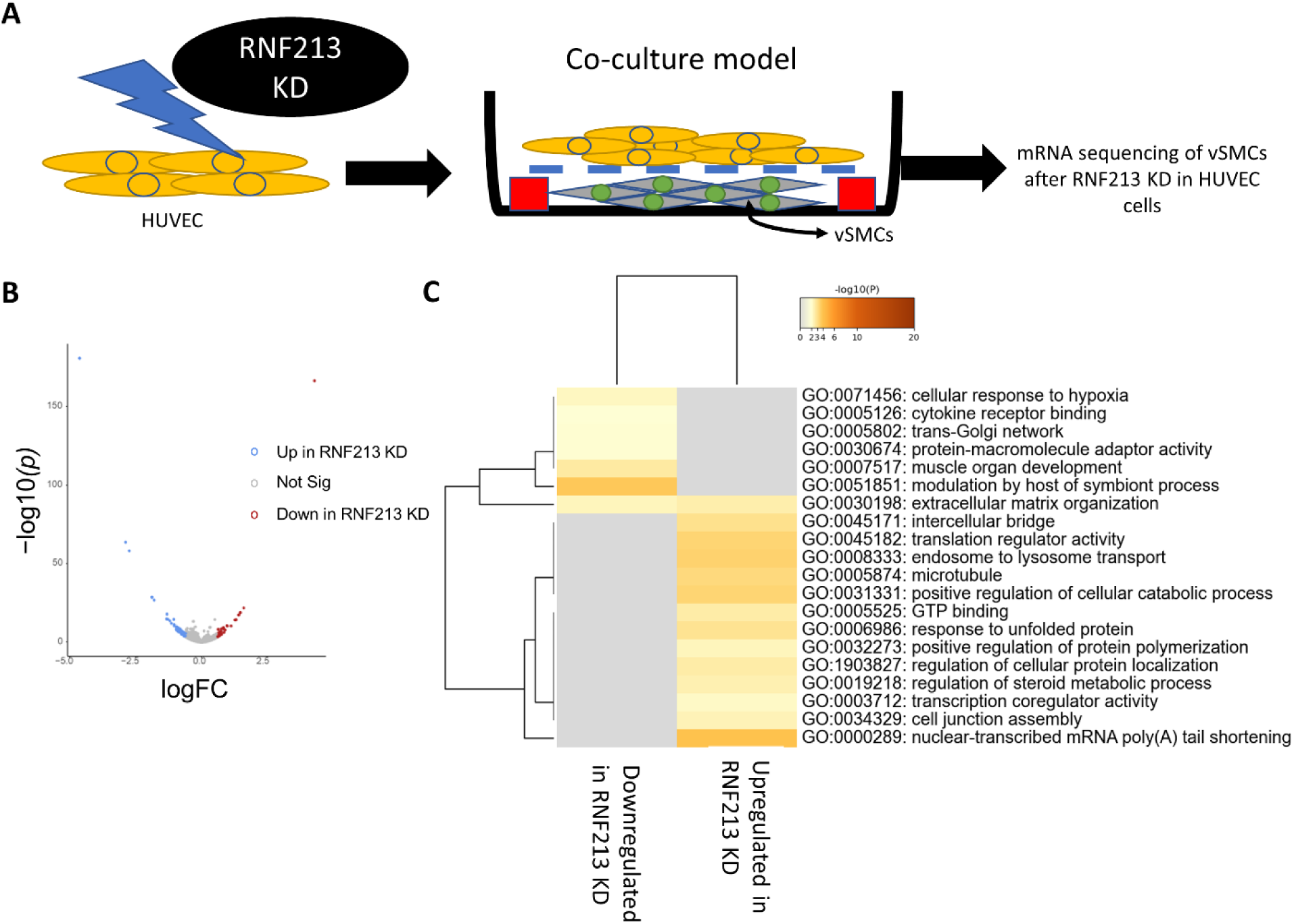
RNF213 KD in HUVEC alters endothelial to vascular smooth muscle cells (vSMCs) communication. A: schematic of the co-culture system. RNF213 was knocked down in HUVEC. Mock transfected or RNF213 KD HUVEC cells were then co-cultured with wild type vSMCs as shown. vSMCs were collected and sequenced. B: Volcano plot of DEGs in vSMCs co-culture model. C: GO analysis (3 terms) for DEGs in vSMCs co-culture model.
➢ **RNF213 function in vascular smooth muscle cells** To complement our data, and to fully grasp the changes brought about by RNF213 loss of function on the vascular system, we performed siRNA KD of RNF213 in vascular smooth muscle cells followed by RNA-seq. siRNA KD led to ≥ 80% downregulation of RNF213 gene expression levels in vSMCs [Figure 8A]. RNA-seq revealed hundreds of DEGs after RNF213 KD [Figure 8B]. GO analysis revealed many overlapping and differentially enriched pathways after RNF213KD in vSMCs [Figure 8C, Supplementary figure 15]. Notably, in the upregulated DEGs, we observed an enrichment of GO biological processes terms that can influence vSMCs migration, such as response to oxidative stress, a known driver of vSMCs proliferation^39^, muscle development, and negative regulation of cell migration [Figure 8C]. Terms related to actin binding, sarcomere, and cytoskeletal organization were also enriched after RNF213 KD, indicating a strong impact on vSMCs function and contractility [Supplementary figure 15]. **Figure 8:**
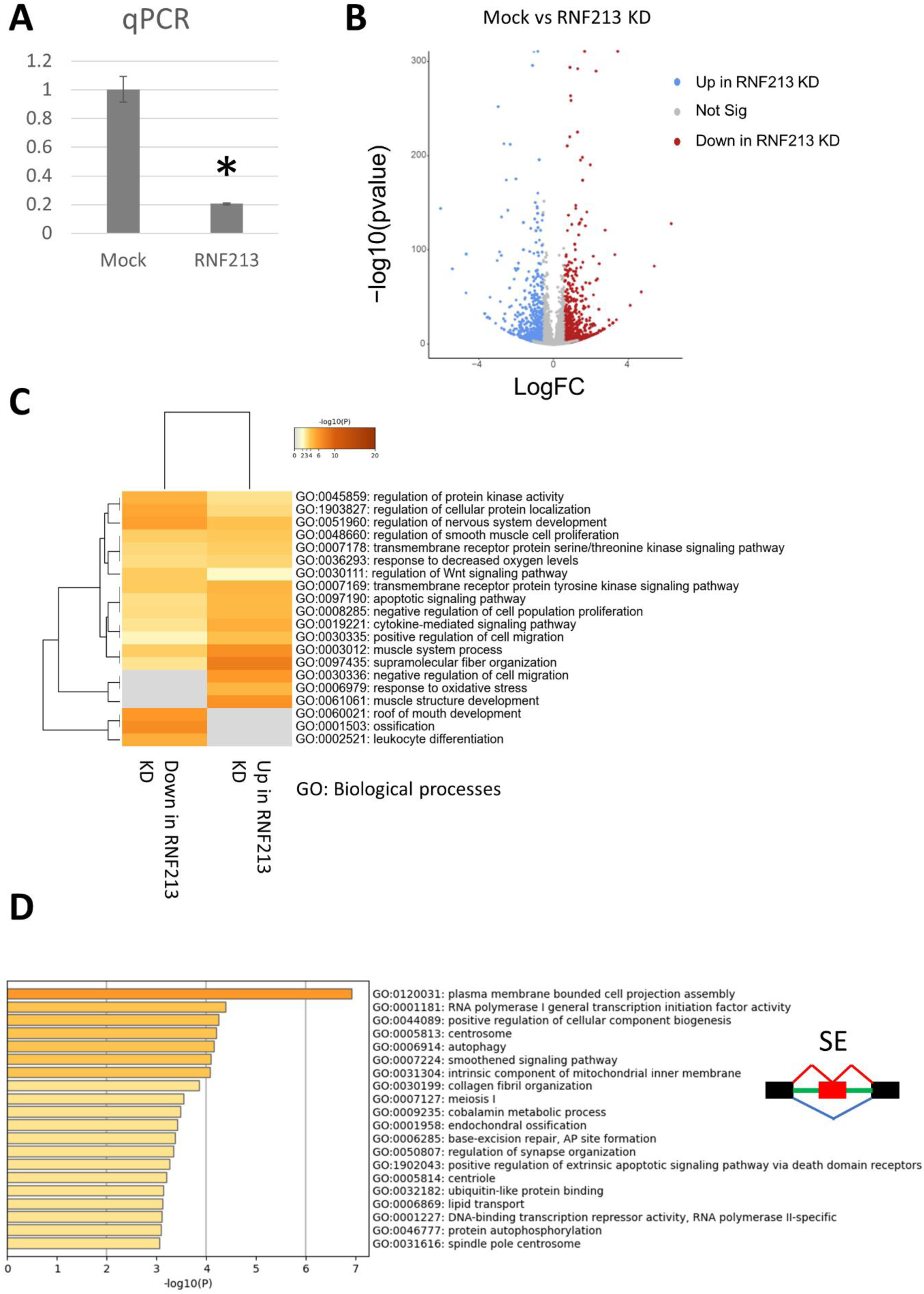
RNF213 KD lead to transcriptional and mRNA splicing alterations in vSMCs. A: confirmation of RNF213 KD using siRNA in vSMCs (Asterisk: statistically significant). B: Volcano plot of DEGs after RNF213 KD in vSMCs. C: GO BP enrichment analysis after RNF213 KD in vSMCs. D: GO enrichment analysis (3 terms) of SE events after RNF213 KD in vSMCs LSV analysis revealed a robust AS program following RNF213 KD in vSMCs, in agreement with the abovementioned observations, with hundreds of events observed in each LSV subtype after RNF213 KD. GO analysis revealed many enriched terms that were specific to each LSV subtype [Figure 8D, supplementary figure 16]. Many of the enriched terms pertained to extracellular matrix, cell migration, cell adhesion, and mitochondrial function. Importantly, the difference of cell processes regulated by different LSV subtypes, and in turn the expected impact on mRNA translation or half-life, indicates a complex interactome that governs the impact of RNF213 loss of function on vSMCs. How these changes observed in vSMCs after RNF213 KD contribute to the medial thinning observed in MMD vessel histology^40^ remain to be fully comprehended.

## Discussion

In this work we provide an in-depth characterization and analysis of the transcriptional and epigenetic impact of RNF213 loss of function in vascular cells. Our aim was to understand how MMD is initiated and how it progresses, as well as how RNF213 mutations and loss of function would promote this pathogenesis. Our data revealed several interesting phenomena. First, we show that RNF213 KD impacts endothelial angiogenesis, migration, and response to inflammation (in the form of LPS). This is mediated via transcriptional reprogramming that impacts cell proliferation and cell surface receptors. Second, we show, for the first time, a strong, albeit indirect, impact of RNF213 on the cellular alternative splicing landscape. Importantly, we show how the pathways regulated by alternative splicing in our dataset can faithfully explain certain phenomena observed after RNF213 KD or in MMD. Third, we show that RNF213 impacts endothelial cell-to-cell communication with leukocytes and with vSMCs. This can explain the links between RNF213 mutations and vascular inflammatory changes, and vessel histology observed in MMD patients. Finally, we show that RNF213 KD leads to transcriptional reprogramming and changes at the alternative splicing level in vSMCs. It’s important to note that the observed transcriptome changes in endothelial cells were very similar to what was observed in microarray analysis of middle cerebral artery specimens from MMD patients^10^. The enriched pathways, at the level of transcriptome in endothelial cells, were also surprisingly similar to dataset obtained from proteome analysis of exosomes in MMD patients^41^. This observation confirms that the observed changes at the exosome level in Wang et al study^41^ are indeed of vascular origin. Overall, our results provide a wealth of information that can be used as blueprint to further study RNF213 gene function and how it regulates vascular biology and its causal links to MMD.

A very interesting aspect of MMD is the exclusive occurrence of stenosis in the carotid terminus, despite being a genetic disease that, theoretically, should impact many vessels in the body. A way to explain this phenomenon is by looking at the blood hemodynamics in MMD patients^26^. However, traditional analysis methods utilizing wall shear stress or other common parameters fail to pinpoint the causality of MMD locality of pathogenesis^24,26^. Interestingly, in a previous unrelated work, our group was able to discover a new type of turbulence in the common and internal carotid arteries in healthy individuals^42^. In the same work, we analyzed various arteries in the human body using mathematical methods, and while the turbulence was observable in all the arteries, and is in fact a property of blood flow as we concluded in our work, the characteristics of turbulence in the carotids were very different, and in fact much more complex, than other vessels^42^. By applying the principles of turbulence to MMD patients, we were able to discover novel hemodynamic phenomena that can explain transient ischemic attacks in early MMD with mild to moderate vessel stenosis^26^. Thus, we propose a link between turbulence hemodynamics and harmonics in the ICA, RNF213 mutations and loss of function, and an altered endothelial response to hemodynamics that initiates ICA terminus stenosis. While some data in our presented analysis does support this hypothesis, such as RNF213 expression levels changing in response to WSS, we cannot test it yet. Unfortunately, the available platforms to study endothelial response to hemodynamics are incapable of faithfully replicating the non-Kolmogorov turbulence observed in the ICA or replicating the specific cardiac harmonics present^24,42^. We hope in the future this theory can be tested with new devices or *in vivo* in animal models of MMD.

While our study gives novel interesting insight into the gene function of RNF213 and its links to MMD pathogenesis, there are some limitations which are inherent to the approach we employed. The main limitation is that we did not test the direct factors contributing to the observed effects. For example, RNF213, as an E3 ubiquitin ligase^7,11,12^, interacts with several proteins to regulate their levels^13^. Downstream of this interaction, changes in RNA binding proteins, transcription factors, or RNA modifying proteins should be responsible for the observed transcriptional and epigenetic changes. Notwithstanding, we believe due to the wealth of data presented, and the exhaustive analysis, analyzing each and every effector would make for a difficult read and will extend the scope of the work beyond what’s desired. Thus, we believe future work should dissect the phenomena presented herein, identify effectors and causal factors, and perform more focused and in-depth analysis to clarify and study the interesting observations presented in our work (in contrast to our systems approach presented in this work).

In conclusion, we propose an improved model for MMD that entails interaction between immune stimulation of ECs, disrupted mechanosensitive responses to blood hemodynamics, and alteration in vSMCs that culminate in the induction of terminal ICA stenosis leading to chronic cerebral ischemia. This stimulates the generation of collateral vessels, which are, in turn, poorly formed due to the effect of RNF213 loss of function inherent in MMD aggravated by yet-to-be-identified immune factors. This model is based on observations made in MMD patients^43,44^, our data, and previously published work^7,13,15^.

## Author contribution

LZ: Conducted experiments. Analyzed Data. Wrote the manuscript. SR: Conception and study design. Conducted experiments. Analyzed data. Wrote the manuscript. Funding. Administration. Supervision. YZ: Conducted experiments. Reviewed the final version of the manuscript. KN: Funding. Administration. Critically reviewed the manuscript. TT: Funding. Administration. Reviewed the final version of the manuscript.

## Data availability

All sequencing data were deposited at the sequence read archive (SRA) project numbers PRJNA779469 and PRJNA782063

