## Supplementary figures for "RNF213 loss of function reshapes vascular transcriptome and spliceosome leading to disrupted angiogenesis and aggravated vascular inflammatory responses"

***Corresponding author**

**# Lead author**

**Corresponding author details:**

Sherif Rashad, M.D, Ph.D.

Kuniyasu Niizuma, MD, PhD.

### Supplementary figures

**Supplementary figure 1:** A: Cluster analysis of enriched GO BP terms after RNF213 KD in HUVEC showing cell division/proliferation cluster to be exclusively enriched in downregulated genes (Supplement to figure 2C). B: GO Molecular functions (MF) enrichment analysis after RNF213 KD in HUVEC.

A

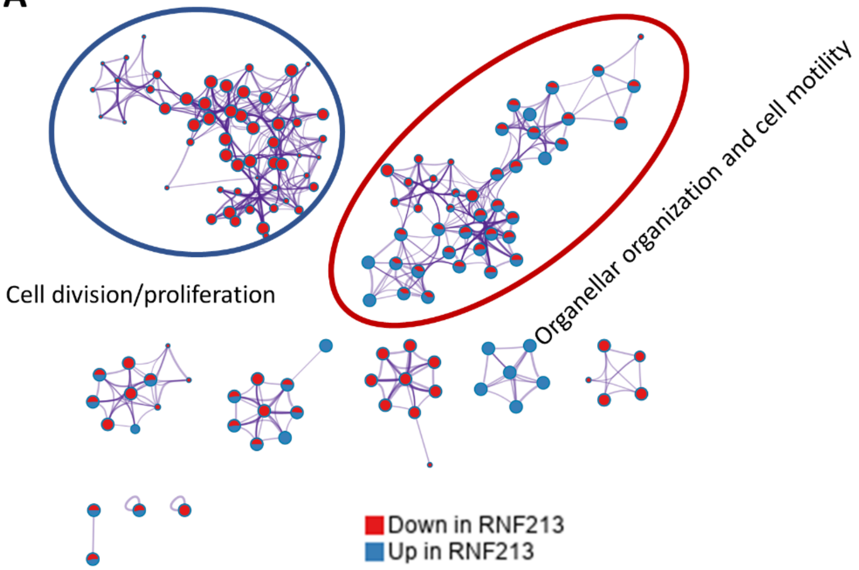


B

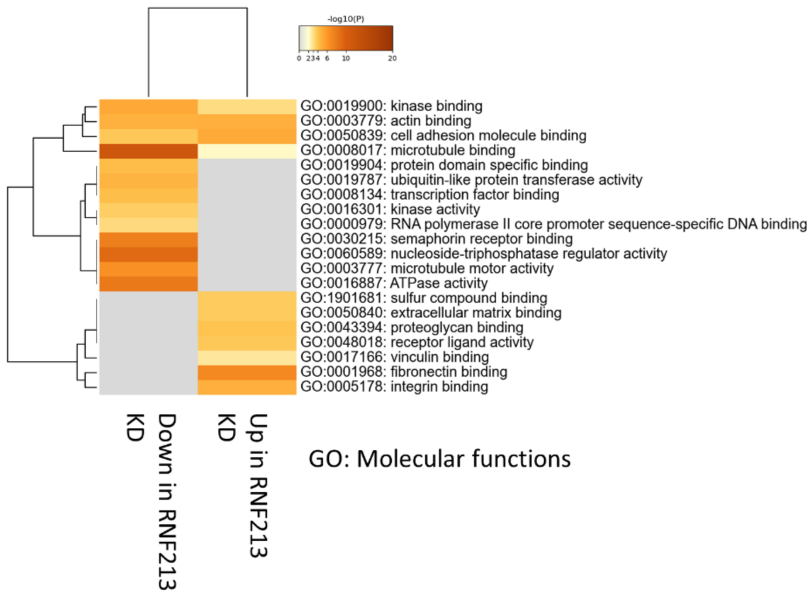

**Supplementary figure 2:** Cluster analysis of GO CC enriched terms revealing two main clusters (A) with differential pattern of enrichment pertaining to each cluster in up and downregulated genes after RNF213 KD in HUVEC (B) (supplement to figure 2D).

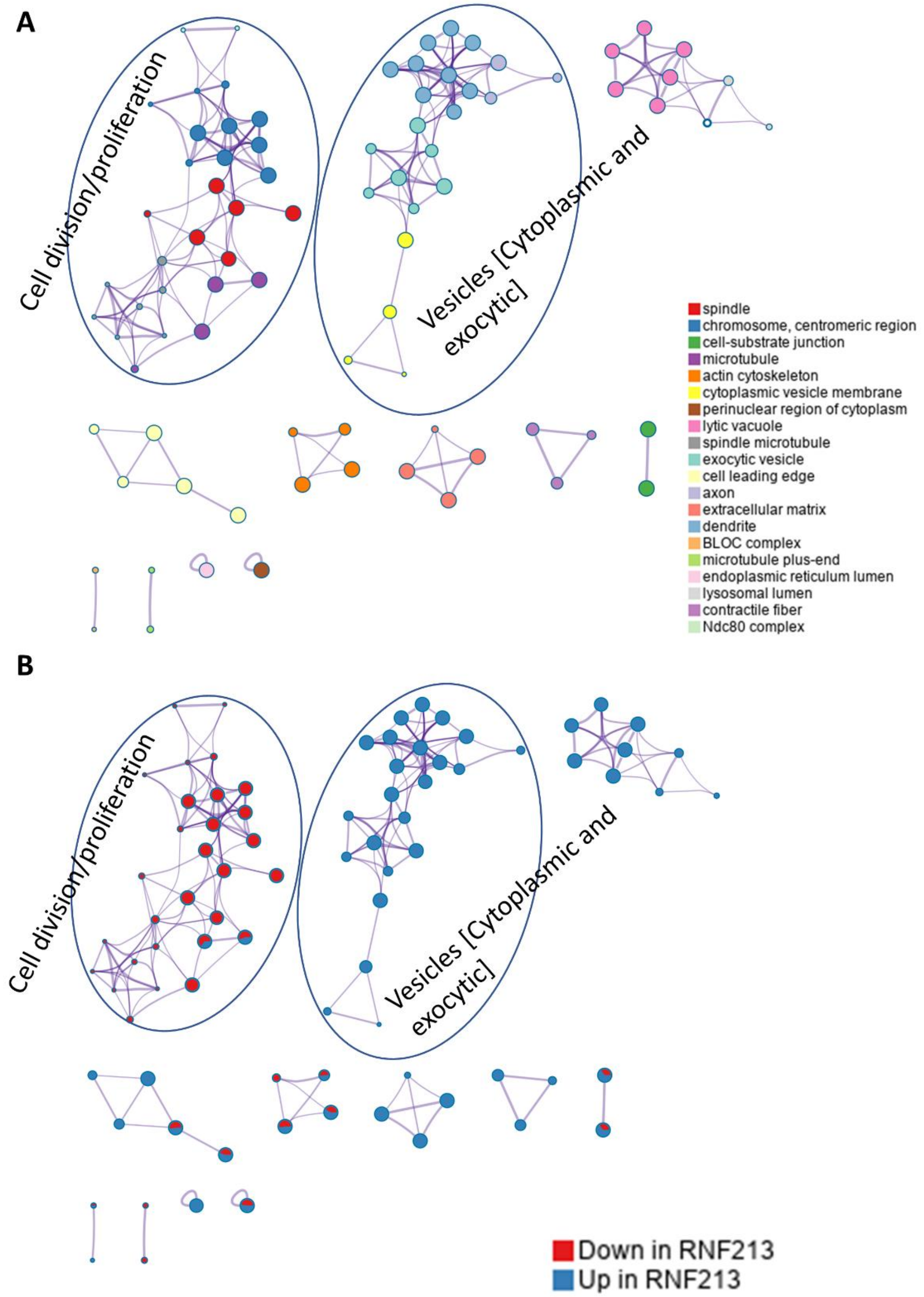

**Supplementary figure 3:** Protein-protein interaction (PPI) network analysis after RNF213 KD in HUVEC. PPI network clusters in downregulated (A) and upregulated (B) genes after RNF213 KD. See supplementary table 1 for MCODE terms.

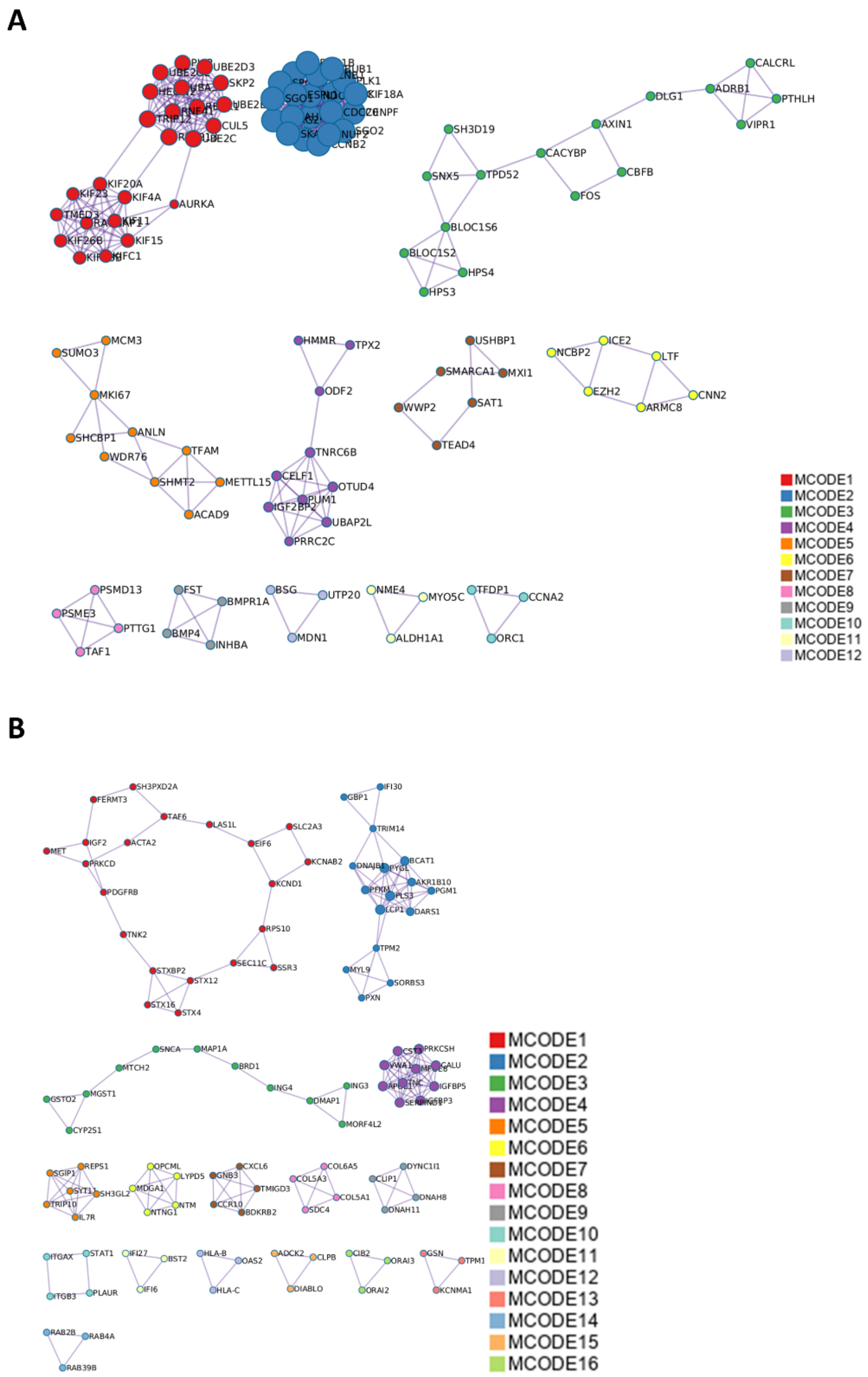

**Supplementary figure 4:** PPI network analysis of all DEGs after RNF213 KD in HUVEC.

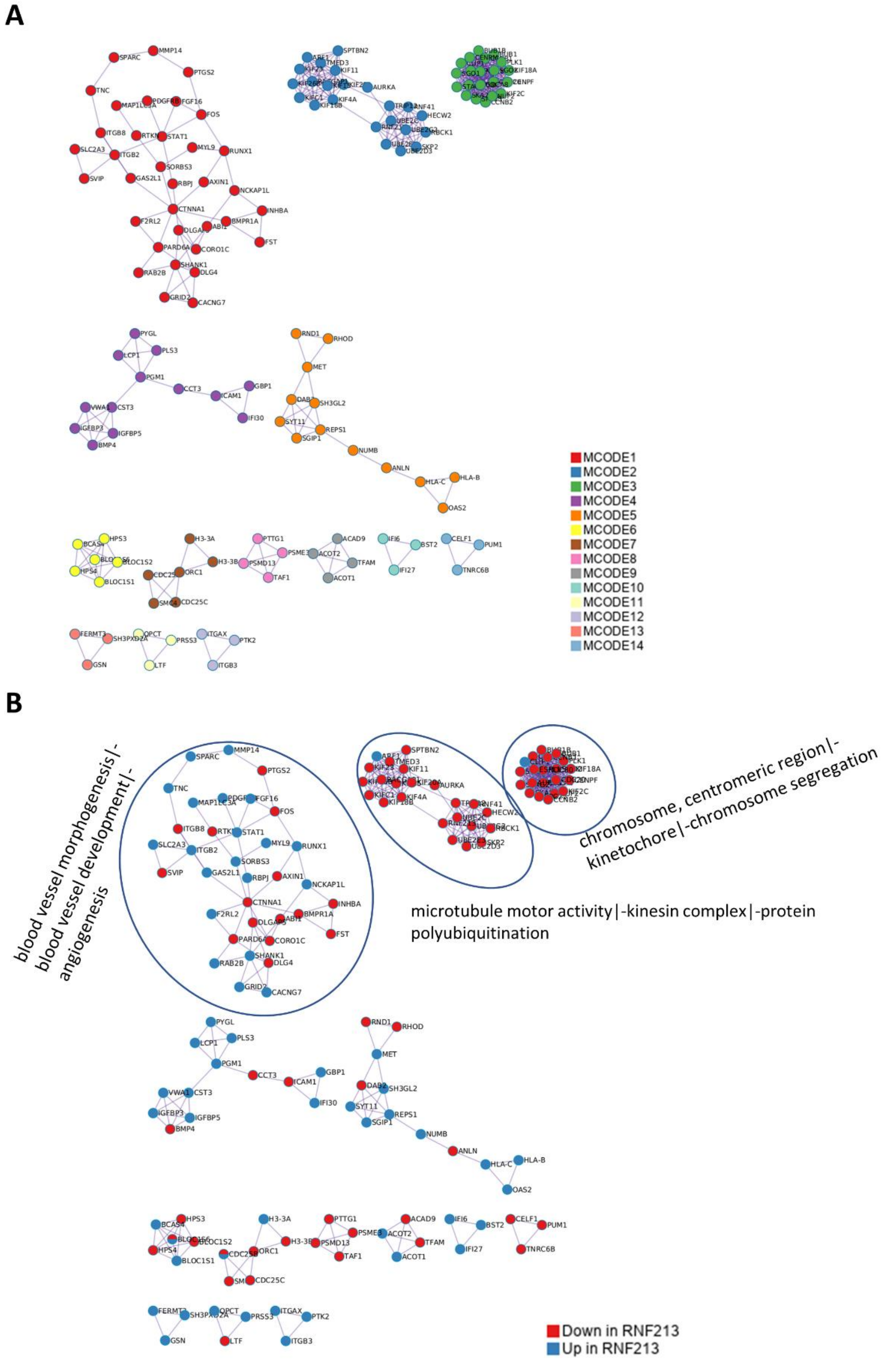

**Supplementary figure 5:** Volcano plots of different LSV subtype events enriched after RNF213 KD in HUVEC (supplement to figure 3)

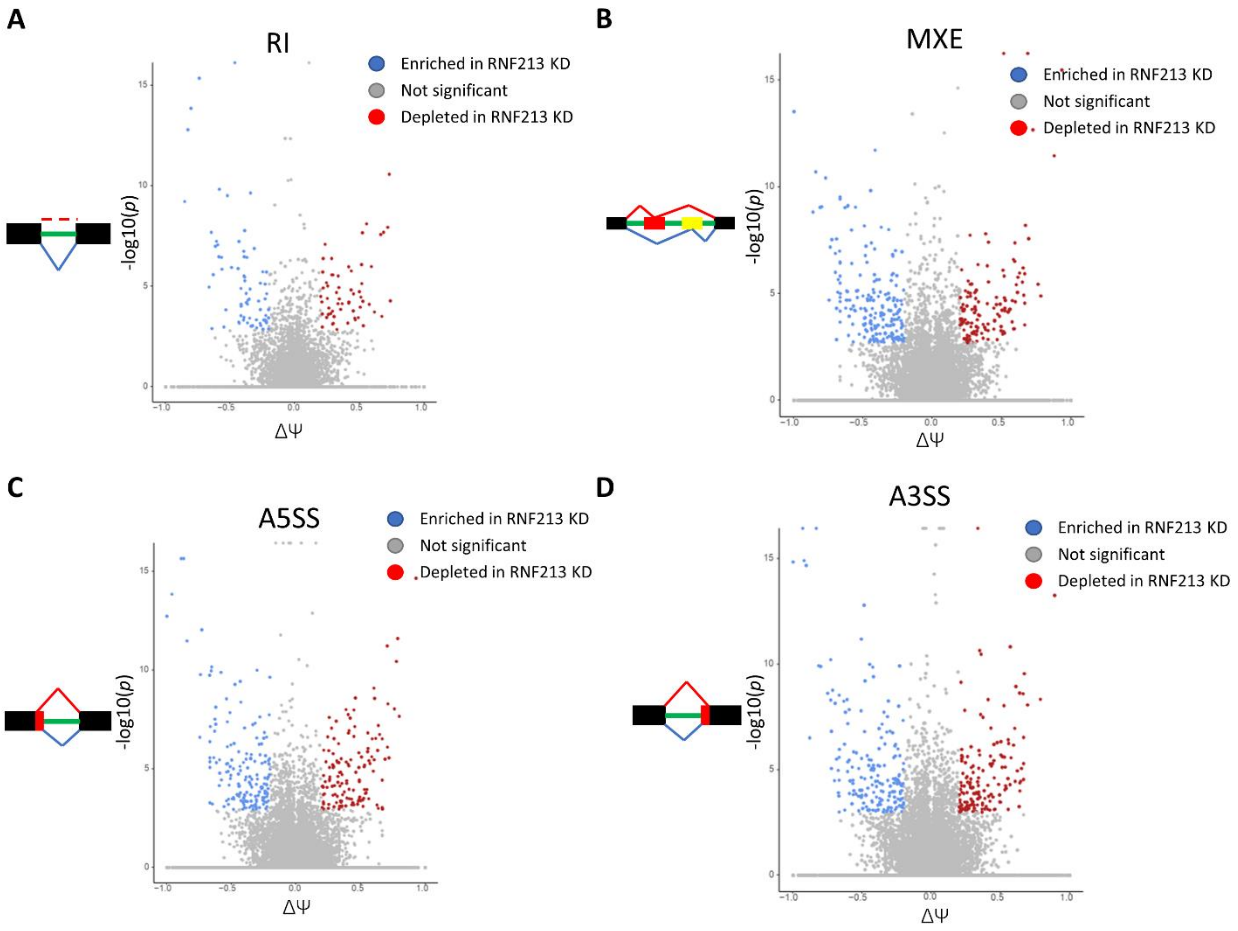

**Supplementary figure 6:** Venn diagrams showing the minimal overlap between alternatively spliced genes and the DEGs after RNF213 KD in HUVEC.

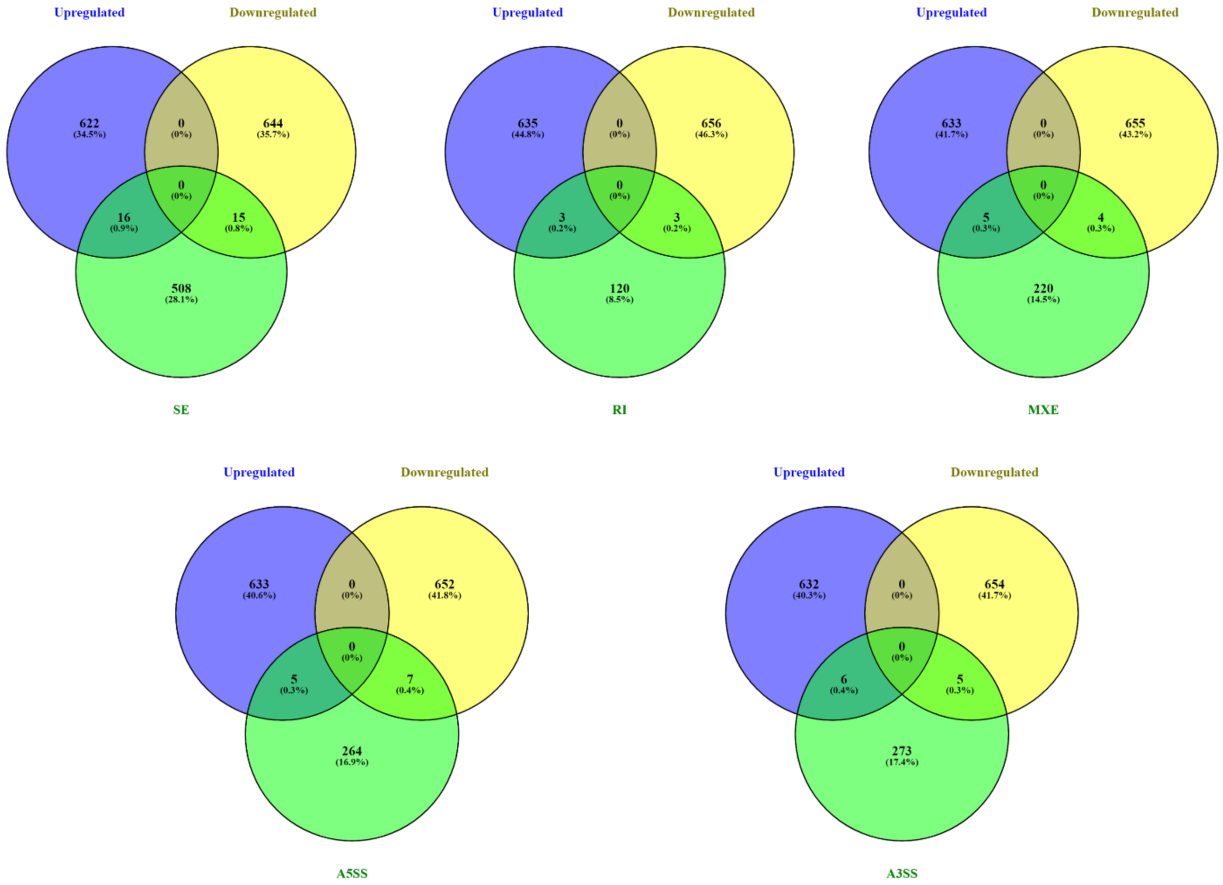

**Supplementary figure 7:** GO (3 terms) enrichment analysis for enriched or depleted LSV events for each LSV subtype (RI not shown).

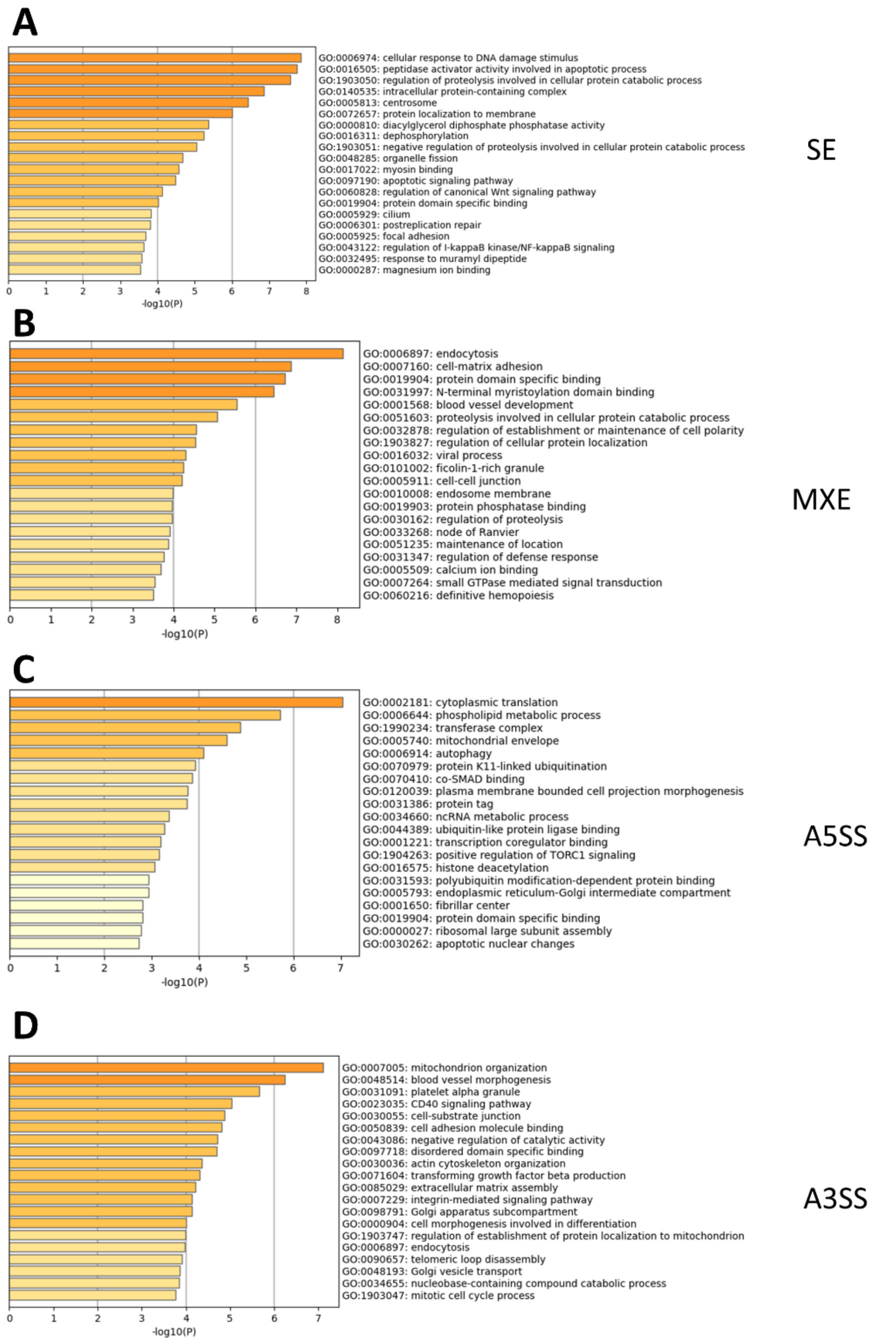

**Supplementary figure 8:** Phase contrast microscope photographs of HUVEC migration after scratch test in the presence or absence of LPS (1ug/ml) in Mock transfected or RNF213 KD groups. The red line represent the cell migrating edge. Distance between the red lines represent the gap across which cells must migrate (Supplement to figure 4B).

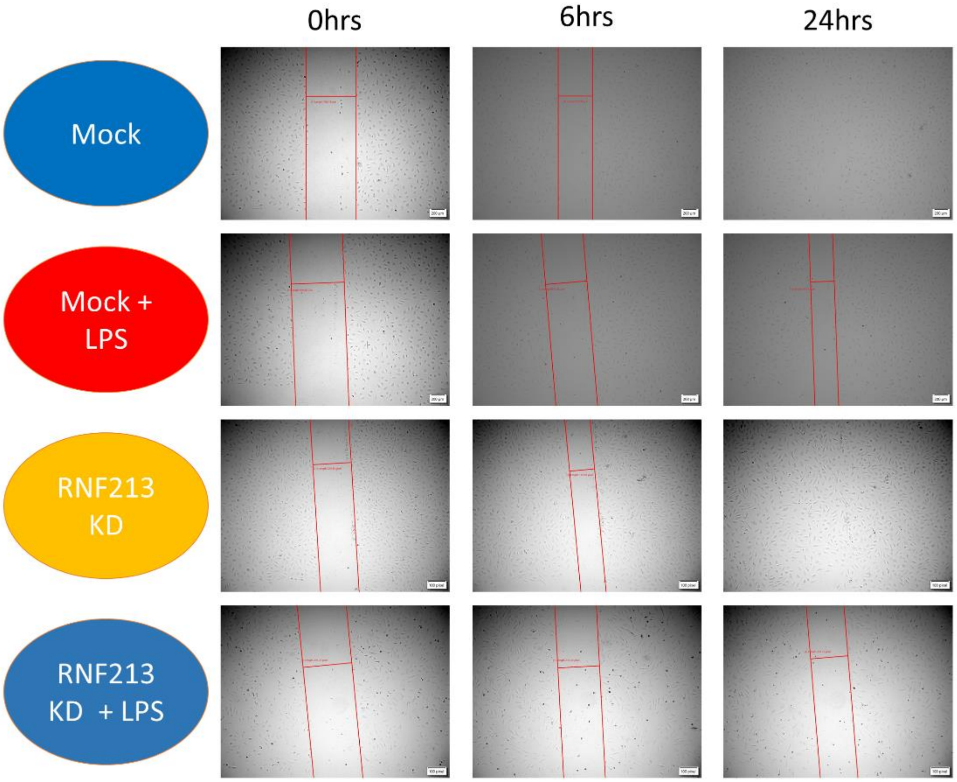

**Supplementary figure 9:** A & B: Venn diagram showing the unique and overlapping DEGs in the Mock or RNF213 groups after LPS treatment. C: GO analysis of uniquely upregulated genes in the RNF213 + LPS group. D: GO analysis of the uniquely downregulated genes in the RNF213 + LPS group.

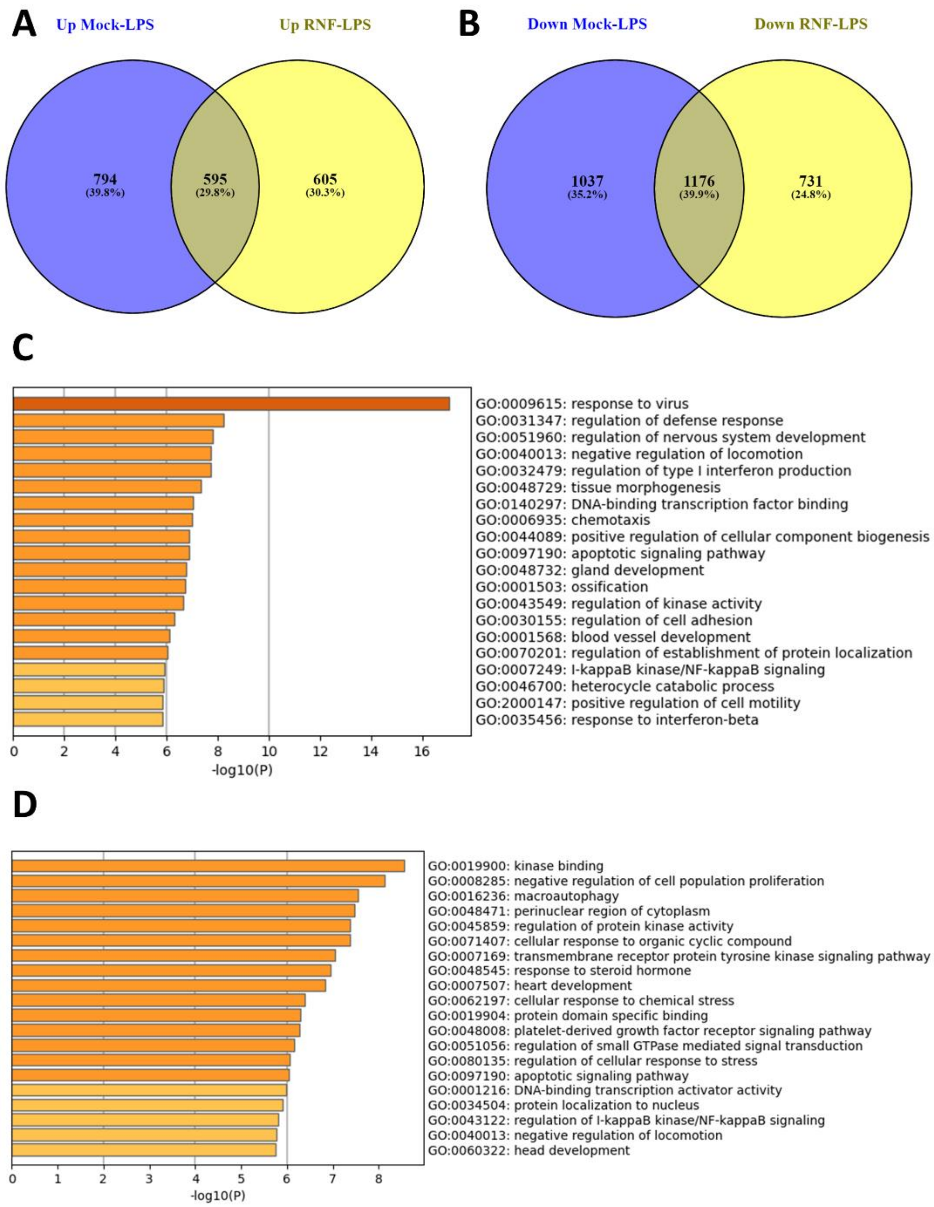

**Supplementary figure 10:** GO analysis of the uniquely upregulate (A) and downregulated (B) genes in the Mock + LPS group.

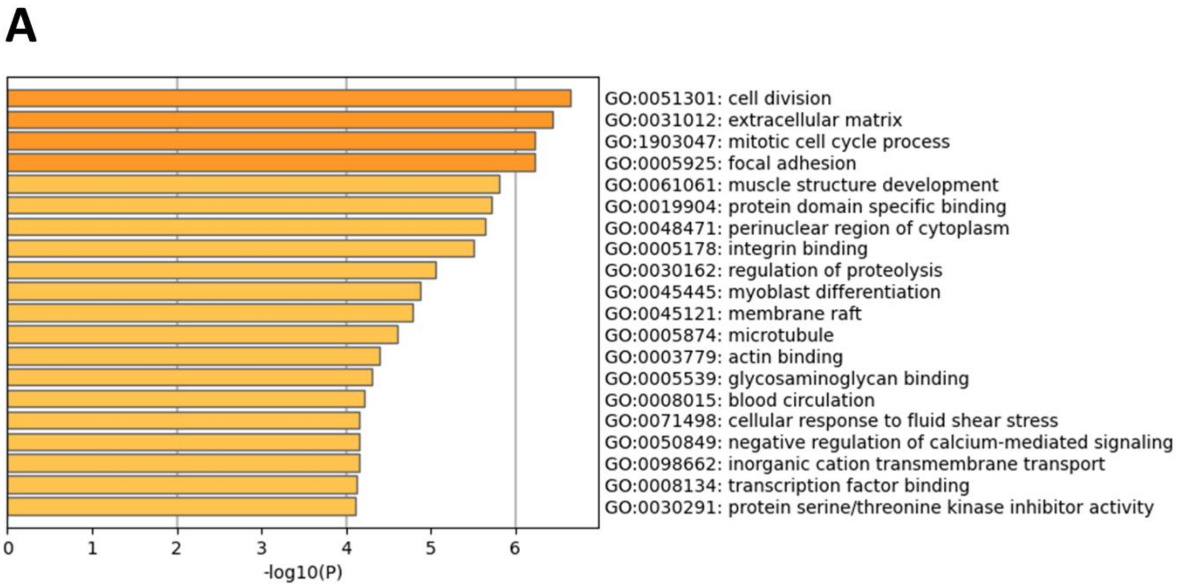


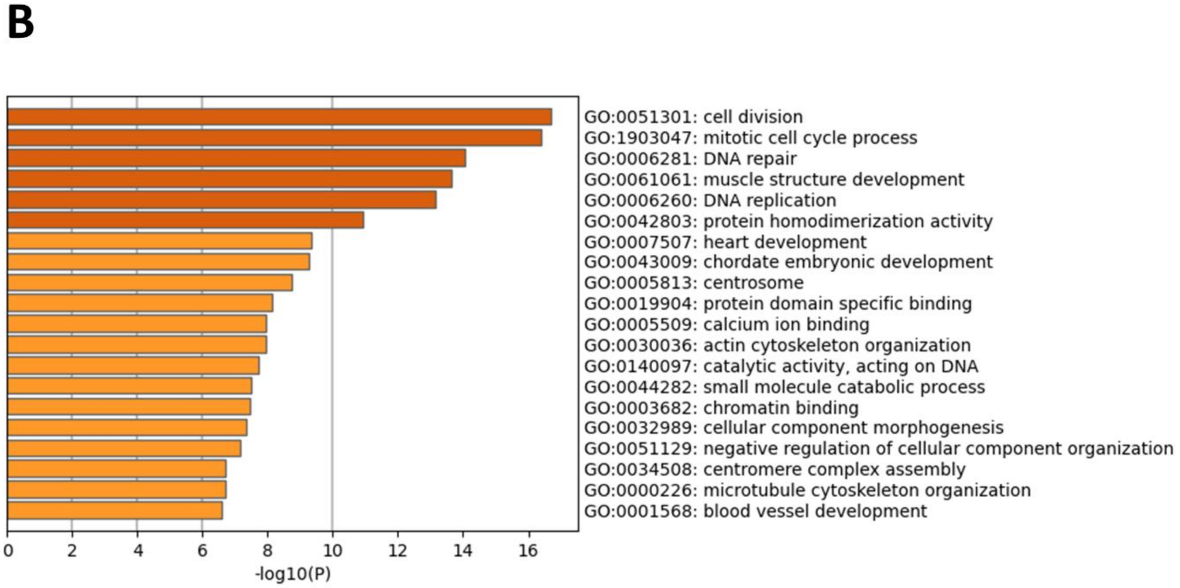

**Supplementary figure 11:** GO BP analysis of DEGs observed when comparing Mock + LPS vs RNF + LPS groups directly (Supplement to figure 5E)

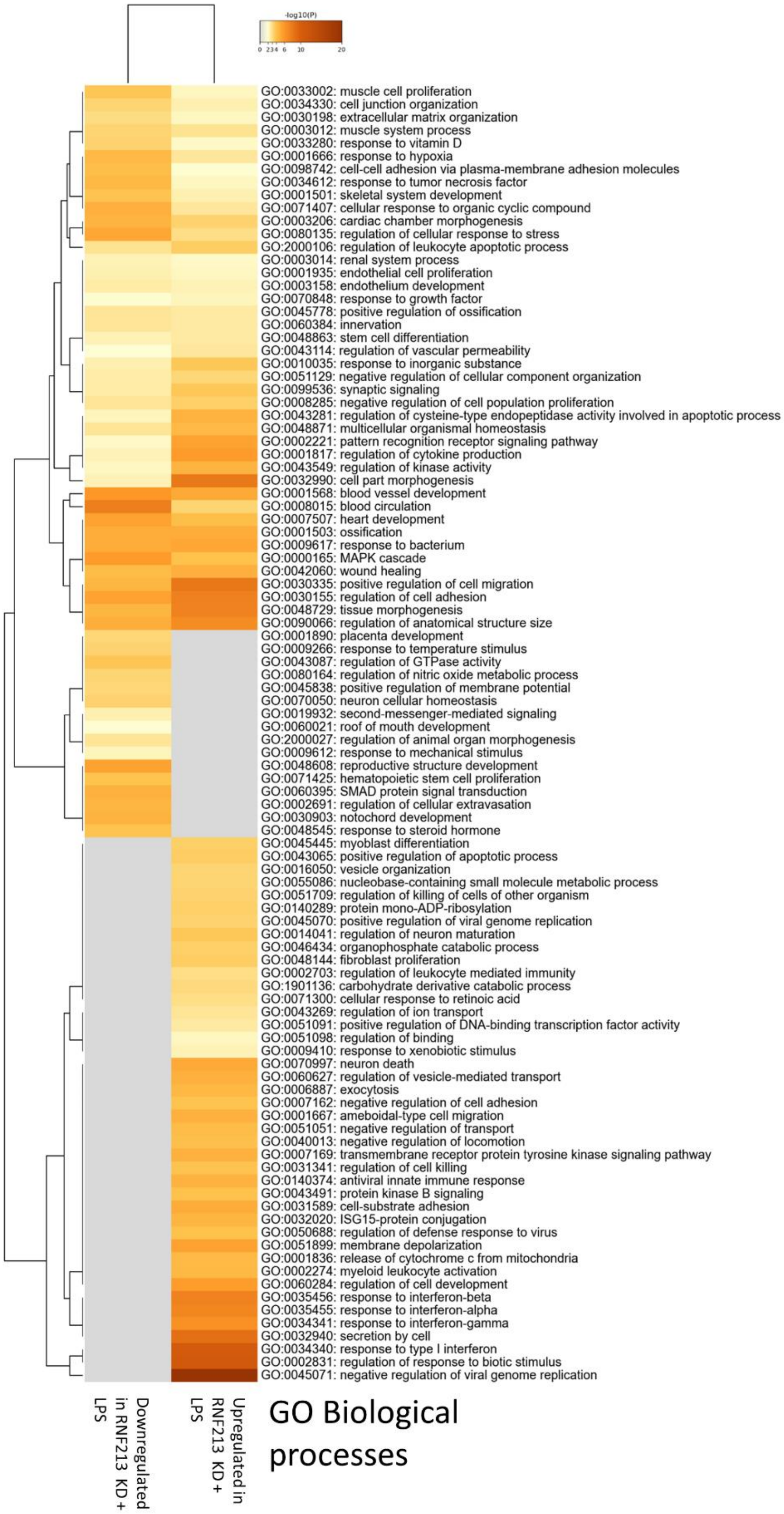

**Supplementary figure 12:** GO MF & CC analysis of DEGs observed when comparing Mock + LPS vs RNF + LPS groups directly (Supplement to figure 5E)

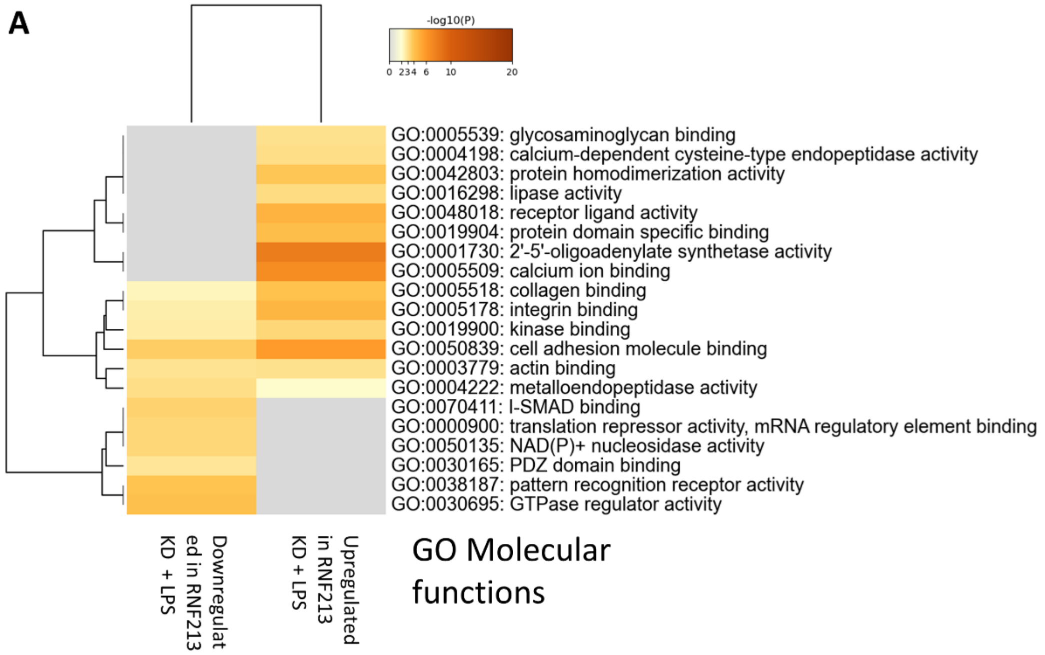


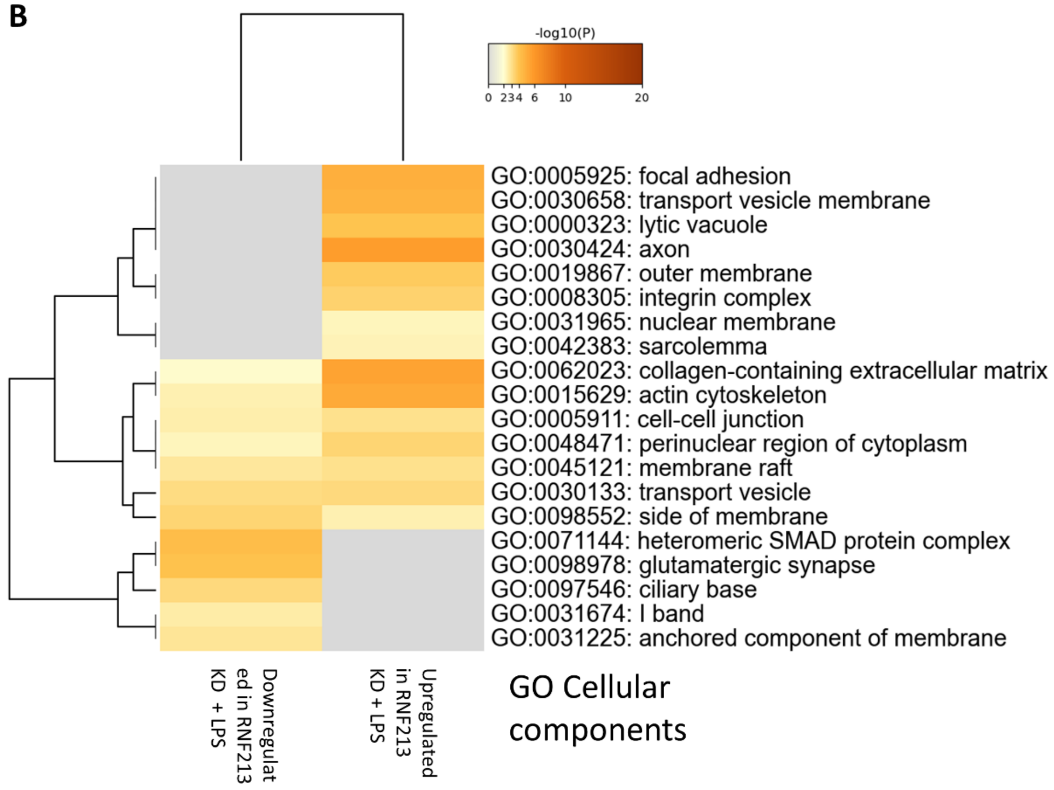

**Supplementary figure 13:** Venn diagrams showing minimal overlap in the observed LSVs when comparing Mock and RNF213 groups after LPS treatment

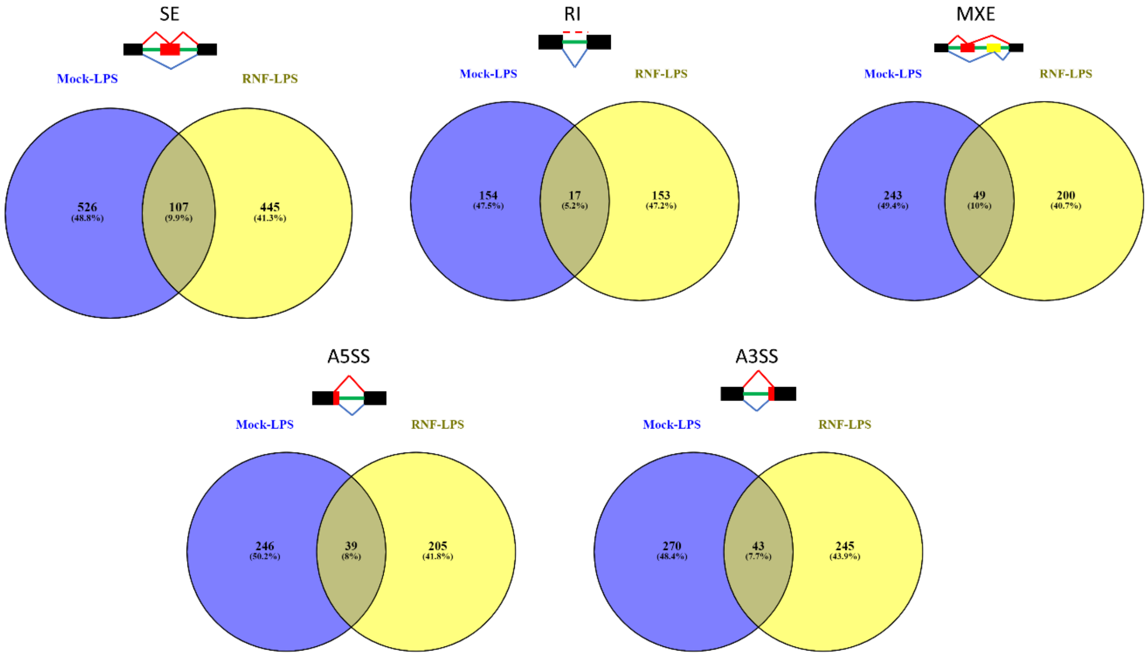

**Supplementary figure 14:** GO analysis (3 terms) for each LSV subtype after LPS treatment in HUVEC (supplement to figure 6C)

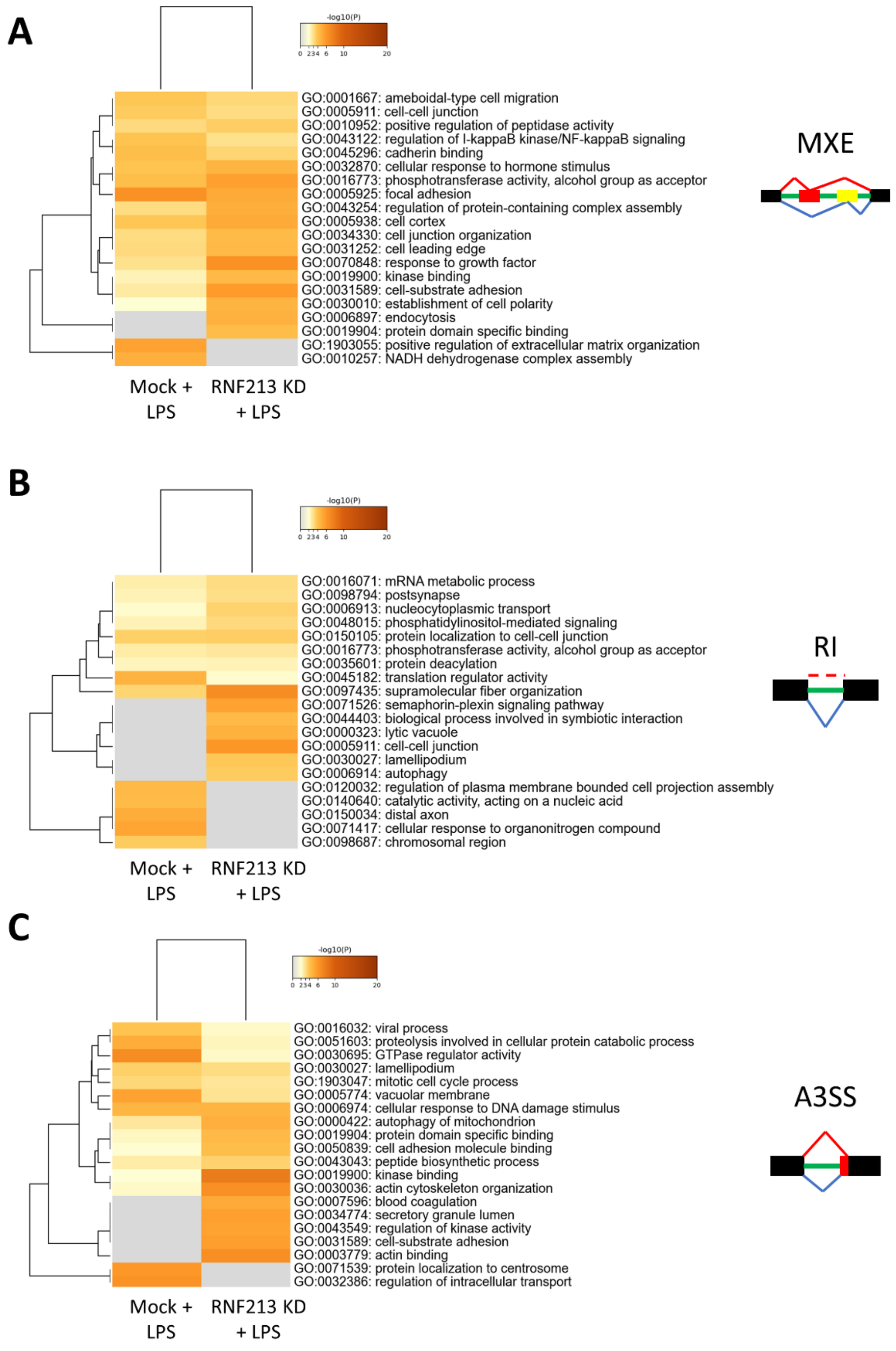

**Supplementary figure 15:** GO MF and CC analysis for DEGs after RNF213 KD in vSMCs (supplement to figure 8C).

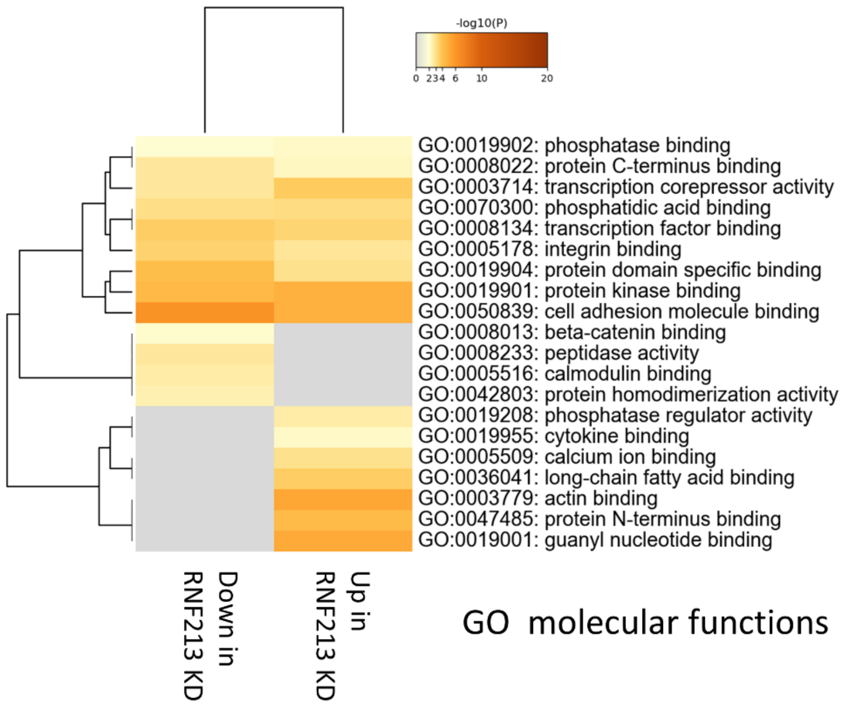


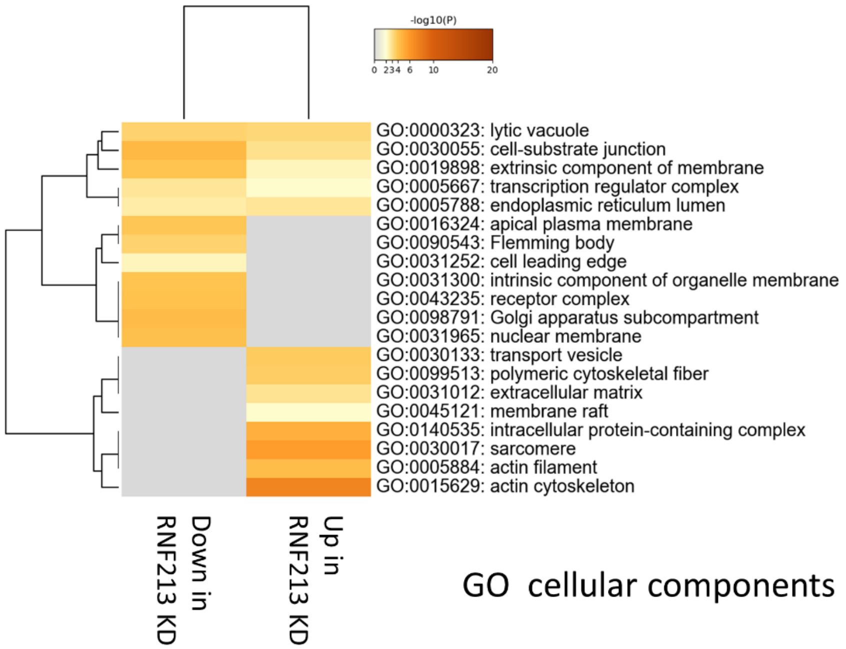

**Supplementary figure 16:** GO (3 terms) enrichment analysis for each LSV subtype after RNF213 KD in vSMCs.

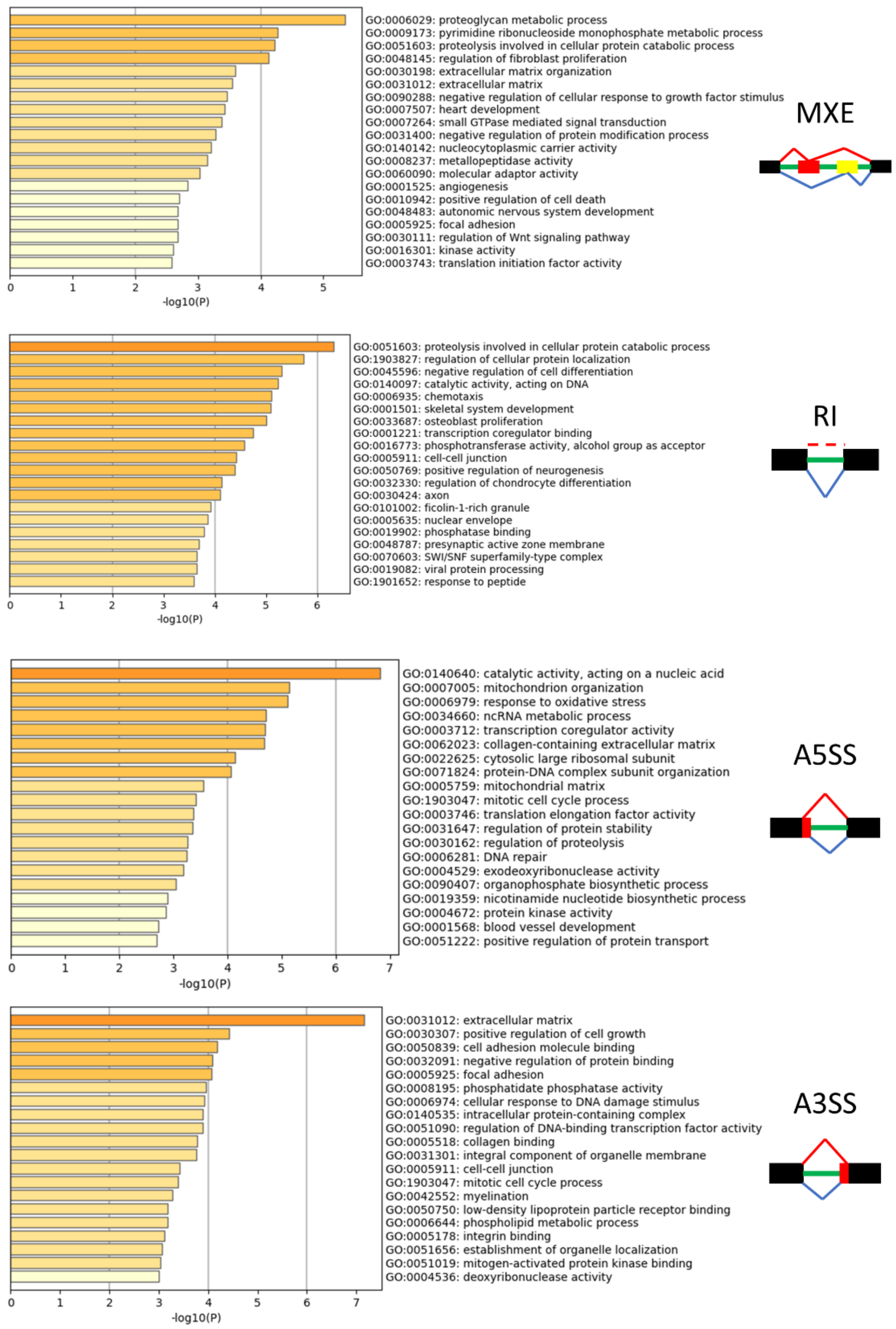
